# Ultrastructural details of mammalian chromosome architecture

**DOI:** 10.1101/639922

**Authors:** Nils Krietenstein, Sameer Abraham, Sergey V. Venev, Nezar Abdennur, Johan Gibcus, Tsung-Han S. Hsieh, Krishna Mohan Parsi, Liyan Yang, René Maehr, Leonid A. Mirny, Job Dekker, Oliver J. Rando

## Abstract

Over the past decade, 3C-related methods, complemented by increasingly detailed microscopic views of the nucleus, have provided unprecedented insights into chromosome folding in vivo. Here, to overcome the resolution limits inherent to the majority of genome-wide chromosome architecture mapping studies, we extend a recently-developed Hi-C variant, Micro-C, to map chromosome architecture at nucleosome resolution in human embryonic stem cells and fibroblasts. Micro-C maps robustly capture well-described features of mammalian chromosome folding including A/B compartment organization, topologically associating domains (TADs), and cis interaction peaks anchored at CTCF binding sites, while also providing a detailed 1-dimensional map of nucleosome positioning and phasing genome-wide. Compared to high-resolution in situ Hi-C, Micro-C exhibits substantially improved signal-to-noise with an order of magnitude greater dynamic range, enabling not only localization of domain boundaries with single-nucleosome accuracy, but also resolving more than 20,000 additional looping interaction peaks in each cell type. Intriguingly, many of these newly-identified peaks are localized along stripe patterns and form transitive grids, consistent with their anchors being pause sites impeding the process of cohesin-dependent loop extrusion. Together, our analyses provide the highest resolution maps of chromosome folding in human cells to date, and provide a valuable resource for studies of chromosome folding mechanisms.

## RESULTS

In eukaryotes, the one-dimensional packaging of chromatin into nucleosomes is understood in great detail, with genome-wide maps of nucleosome positions and composition available for scores of organisms (*1*). Although our understanding of three-dimensional folding of the genome somewhat lags the 1D picture, over the past decade, a wide variety of technical approaches have provided impressively concordant views of chromosome architecture in vivo, as for example chromosome compartments and TAD organization are readily captured using microscopy-based methods (*2, 3*), chromosome conformation capture (3C)-based methods (*4-6*), and orthogonal methods such as genome-architecture mapping (*7*). The majority of genome-wide studies of chromosome folding utilize 3C-based proximity ligation methods (*8*), in which genomic loci in physical proximity are crosslinked to one another, chromatin is fragmented using restriction enzymes, and interacting genomic loci are then identified following ligation and paired-end deep sequencing.

The fundamental resolution of chromosome conformation capture is largely defined by the size and uniformity of chromatin fragmentation, with the coarseness of genome fragmentation providing a lower limit to the resolution of 3C experiments. To approach the maximum practical resolution to 3C methods, which is set by the ubiquitous packaging of the genome into repeating nucleoprotein subunits known as nucleosomes (*9*), we recently developed Micro-C, a Hi-C protocol in which chromatin is fragmented to mononucleosomes using micrococcal nuclease (MNase), thereby increasing both fragment density as well as uniformity of spacing (*10, 11*). Although a substantially updated version of this protocol with improved signal-to-noise was originally named Micro-C XL (*11*), given that this protocol completely supersedes the original one we will simply refer to the updated protocol as Micro-C throughout. The Micro-C protocols were developed in budding and fission yeast, and provide insight into chromosome folding at scales ranging from single nucleosomes to the entire genome.

To extend our Micro-C analysis of chromosome folding from the relatively simple yeast genome to the more complex chromosomal organization seen in mammals, we generated deeply-sequenced Micro-C datasets (∼2.6-4.4 billion uniquely mapped reads per sample, ∼150X coverage per nucleosome – **Table S1**) for two well-studied human cell types: pluripotent human embryonic stem cells (H1-ESC) and differentiated human foreskin fibroblasts (HFFc6). To benchmark our approach, we also generated Hi-C datasets for these two cell types in parallel, following the *in situ* Hi-C protocol with a 4 bp cutter (DpnII) (*12*). The increased signal-to-noise of Micro-C libraries was already apparent at the initial data processing step, where Micro-C libraries yielded a far lower fraction of trans-chromosomal interaction pairs (13.5% for Micro-C vs. 51.8% for Hi-C for ESCs; 15.2% vs. 30.7% for HFFs), which are strongly enriched for random ligation products, thus noise. To visually compare Hi-C and Micro-C views of human cells, we plotted contact heatmaps for ESCs and HFFs in **Fig. 1A-B** at four scales arranged from chromosome-scale (left) to single-gene resolution (right). Overall, Micro-C and Hi-C maps reveal the same major classes of patterns that illuminate various features of chromosome folding: at lower resolutions (e.g, ∼100 kb bins) the coarse checkerboard pattern reflects the compartmentalization of active and inactive chromatin, while zooming to higher resolutions (e.g. <10 kb bins) reveals finer compartmental segmentation, TADs, and “off-diagonal” interaction peaks thought to result from a high frequency of CTCF-anchored long-range looping interactions.

**Fig. 1.**
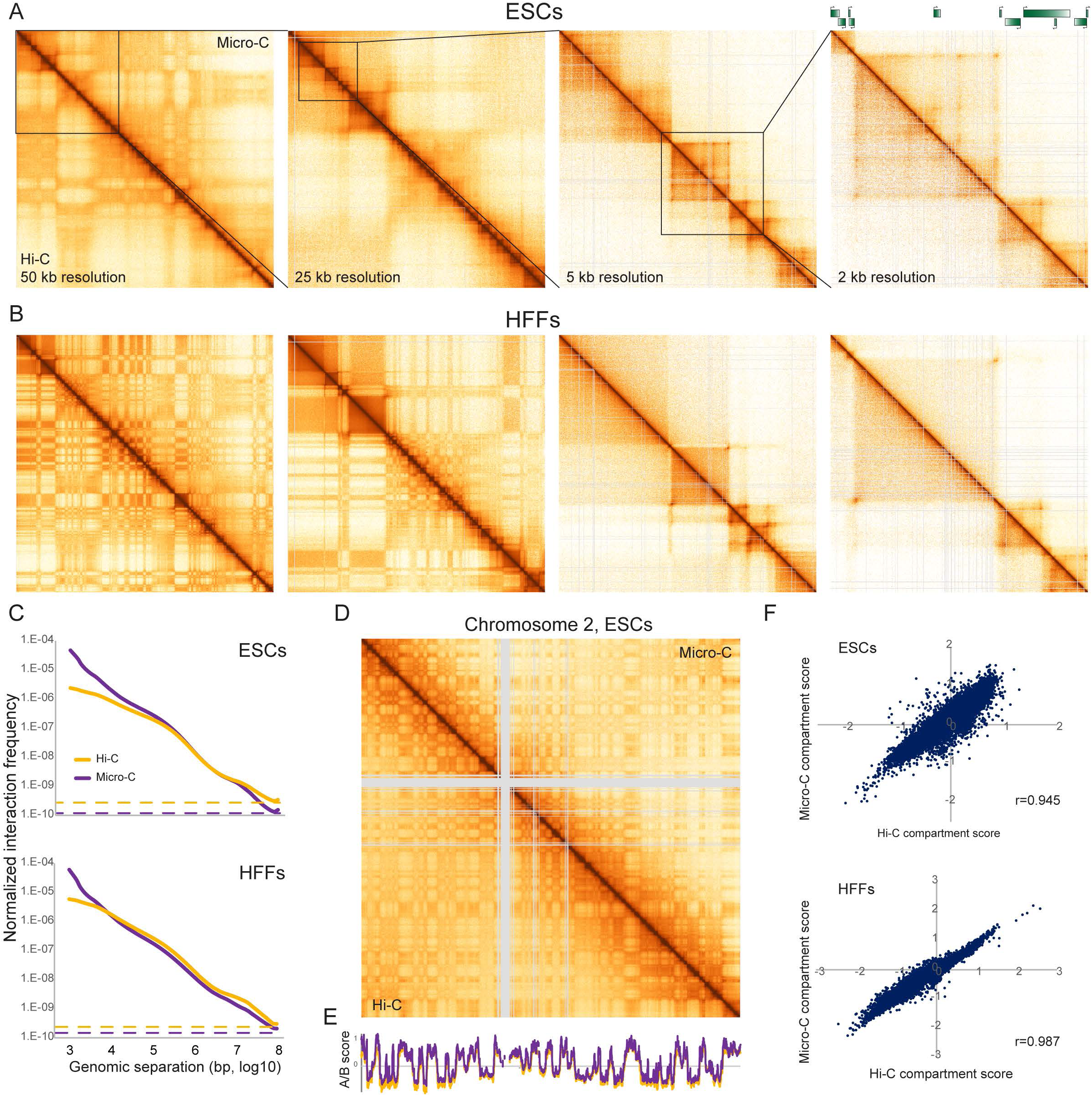
Micro-C of human pluripotent and differentiated cell types recovers coarse features of chromosome folding. (A-B) Chromosome contact maps for four successive zoom-ins across human chromosome 3, for H1 ESCs (A) and HFFs (B). From left to right: Chr3:0-55MB, 0-22 MB, 1-6 MB, 3-5 MB. Each panel show Micro-C data above the diagonal, with Hi-C data for the same cell line below the diagonal. (C) Interaction frequency is plotted for Micro-C and Hi-C on the y axis as a function of genomic distance between interacting fragments (x axis), for ESCs and HFFs. Both axes are in log10 scale. In both cases, dotted lines show the genome-wide average interaction frequency between loci located on different chromosomes, an estimate of nonspecific dataset noise. See also **fig. S1**. (D) Micro-C robustly captures A/B compartment organization. Heatmap shows Hi-C and Micro-C interaction maps (binned at 100 kb resolution) for Chromosome 2 in ESCs, illustrating the nearly identical A/B “checkerboard” pattern captured by both methods. See also **fig. S3**. (E) Eigenvector scores for chromosome 2 in ESCs compared for the Hi-C dataset (orange) vs. the Micro-C dataset (purple). (F) Eigenvector scores are globally correlated between Hi-C and Micro-C maps. Scatterplots show A/B scores for 100 kb genomic tiles in Hi-C (x axis) vs. Micro-C (y axis) maps of ESCs and HFFs.

To broadly survey the performance of Micro-C relative to Hi-C methods, we examined the scaling of contact frequency P(s) as a function of the distance between two genomic loci. Comparing Hi-C and Micro-C, we find a similar decay in interactions with increasing distance for length scales from ∼20,000 bp to 1 Mb (**Fig. 1C**). However, Micro-C exhibits an increased dynamic range of contact frequency, with improvements at both small and large genomic separations. At large separations (>10 Mb), where contacts are expected to be very rare, P(s) curves flatten to approximately the levels of interchromosomal contact frequency (**Fig. 1C**), which is 2 to 4-fold lower in Micro-C than in Hi-C. The average reduction in both interchromosomal and extremely long-range intrachromosomal contact frequencies observed in Micro-C is likely due to lowering of the “noise floor” of artifactual contacts (random ligations between nucleosomes) (*11*). At shorter scales, Micro-C consistently recovers a higher fraction of close-range “near-diagonal” contacts, providing greater coverage of short-range chromosome folding behaviors at the 1-100 nucleosome scale (**fig. S1**). This length scale includes the 1-5 nucleosome scale that is inaccessible to traditional Hi-C due to the longer average fragment size (∼256 bp on average for complete DpnII digestion, with actual fragment sizes closer to ∼1 kb in practice due to partial digestion) and the heterogeneity of fragment lengths resulting from uneven spacing of restriction sites (**fig. S2**). Importantly, this length scale has the potential to provide information about the arrangement and interactions of nucleosomal arrays and about chromatin fiber structure (*13*).

At low and intermediate resolution, Micro-C data recapitulates the chromosome organization seen in Hi-C maps. For example, A/B compartment calls are strongly correlated (r=0.945 and 0.967 for ESCs and HFFs, respectively, at 100 kb resolution) between Micro-C and Hi-C maps (**Fig. 1D-F, fig. S3**). Although many compartments encompass relatively large (∼1 Mb) genomic intervals, consistent with prior reports (*14*) we also find many examples of extremely fine single-gene scale compartments (exhibiting the “plaid” pattern typical of compartments and thus distinct from TADs) in both cell types (**fig. S3A**). Intriguingly, we find a dramatic increase in compartment “strength” in fibroblasts in both datasets (**fig. S3B**), presumably reflecting both the unusually short G1 phase of the ES cell cycle, as well as increased regulatory specialization observed as cells differentiate. While Micro-C and Hi-C maps therefore provide highly concordant views at the coarsest level of genomic organization, they are clearly distinct at higher resolution, as for example a number of interaction peaks (“dots”) are apparent in the Micro-C dataset that are absent in Hi-C, as can be seen in the right panels of **Figs. 1A-B**.

One unique feature of the Micro-C protocol, compared to restriction enzyme-based 3C methods, is that all interacting chromatin fragments are mononucleosomes. As a result, ignoring read pairs and treating the dataset as a single-end MNase-Seq dataset provides a genome-wide map of nucleosomes for a given cell type “for free” (with the possible caveat that it will miss nucleosomes that are crosslinking-or ligation-resistant). To illustrate this feature and to explore the 1-dimensional landscape of ESC and HFF chromatin, we extracted single-end sequencing data from our Micro-C datasets. Consistent with genome-wide analyses of nucleosome positioning across a wide range of species (*15*), aligning the Micro-C nucleosome mapping dataset at promoters confirms the expected nucleosome depletion at active promoters, with nucleosome depletion scaling with transcription rate (**Fig. 2A**). Similarly, we confirm the role for CTCF in establishing local nucleosome patterning (*16-18*) (**Fig. 2B-C**), again validating the utility of the unpaired single-end dataset as a high-quality nucleosome mapping dataset and providing a valuable ultra-high depth resource for future studies investigating nucleosome positioning in these widely used cell types.

**Figure 2.**
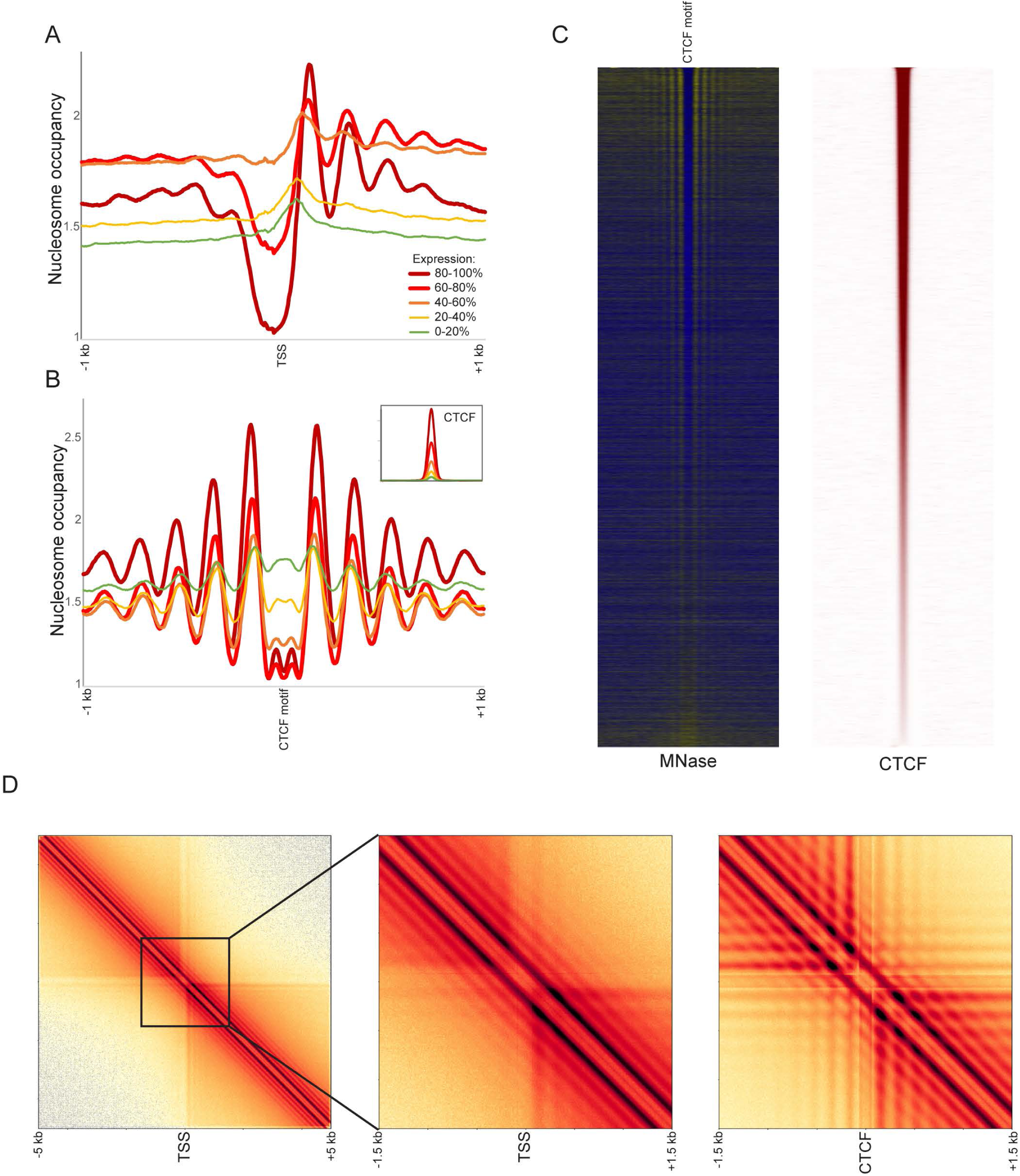
Nucleosome-resolution views of chromatin organization. (A-C) Micro-C recovers nucleosome-resolution chromatin organization. In these panels, read pairing information was discarded, and single-end Micro-C reads (representing nucleosome ends) were shifted 73 bp to the nucleosome dyad axis. (A-B) show nucleosome occupancy profiles aligned according to TSSs (A) or CTCF binding sites (B) and averaged according to quintiles of transcription rate or CTCF occupancy. Panel (C) shows nucleosome occupancy and CTCF ChIP enrichment (*29*) at genes sorted from high (top) to low (bottom) CTCF occupancy. (D) Nucleosome resolution contact maps surrounding TSSs (left two panels) or CTCF binding sites (right panel). Promoters and CTCF sites act as boundaries between contact domains, as seen in the clearing of contacts in the upper right and lower left quadrants. Also apparent at this resolution is the nucleosome phasing surrounding these regulatory elements, manifest as a grid-like structure superimposed on the contact maps.

Including the information contained in the proximity ligations between nucleosomes allows us to use Micro-C to its full potential and capture local features of the chromatin fiber that are beyond the fundamental resolution of Hi-C. Given the key roles for CTCF and promoters in organizing local 1D chromatin, we started by exploring the local nucleosome-nucleosome interactions around TSSs and CTCF binding sites. Averaged read-level interaction maps centered at these elements (**Fig. 2D**) produce a characteristic pattern of accumulation (“inverted egg-carton”), with two distinct banding patterns: (i) a series of horizontal/vertical bands decaying from the center, reflecting the coherent positioning of individual nucleosomes in the vicinity of a DNA-bound factor and (ii) a series of bands parallel to and decaying away from the main diagonal, reflecting a coherence in the interactions among neighboring nucleosomes throughout the fiber. Importantly, this diagonal banding is locus-independent, as manifested in the oscillations of the global interaction decay curves (**fig. S1**), indicating that nucleosomes are organized into regularly spaced arrays genome-wide, regardless of how coherent the positioning of such arrays may be from cell to cell at any given locus. In other words, although the +6 nucleosome in a coding region may exhibit “fuzzy” positioning, occupying different positions in different cells in a population, the adjacent nucleosomes are consistently well-positioned relative to this nucleosome.

To further explore local chromatin fiber behavior in our Micro-C maps, we focused on the behavior of interaction decay curves at short (1-100 nucleosome) distances (**fig. S1**). The behavior of the contact frequency decay at short distances allows us to discount a solenoid-like organization as we find no enrichment of N/N+5 or N/N+6 ligation products. In contrast, we find a subtle signature for a given nucleosome to interact with similar efficiency with adjacent pairs of distal nucleosomes (eg N+2 and N+3, or N+4 and N+5 – see **fig. S1D**). Thus, our data support the model that small (∼3-10) “clutches” of nucleosomes are organized in a 2-start or zig-zag orientation to form short tri-or tetranucleosome zig-zag motif (*10, 13, 19-24*). Although the precise underlying structure of the chromatin fiber cannot be directly extrapolated from our data, the interaction decay curves provide strong experimental constraints to test theoretical models.

Next, we turn to the organization of domains and boundaries in human cells. TADs are chromatin domains thought to emerge as the result of continuous ATP-dependent extrusion of chromatin loops by cohesin. Simulations have shown that the stopping of cohesin, e.g. by DNA-bound CTCF molecules, can create TAD boundaries, as well as other patterns of contact enrichment observed in Hi-C maps (*25, 26*). These signatures include a “square” or “box” on the diagonal of elevated contact frequencies delimited by sharp boundaries (*4, 6*), “stripes” along the box edges (*25*) and “dots” at their far corners (*27*). Stripes and dots are attributed to the stalling of extruded loops at one or two inward-oriented CTCF binding sites, respectively.

To leverage the enhanced resolution afforded by Micro-C to precisely identify genomic correlates of boundary activity (**Fig. 3A**), we defined chromatin interaction boundaries by scanning the genome for local minima in the number of crossing interactions (normalized relative to the local interaction frequency at the same distance (*28*)), i.e. insulation strength. Boundary calls were robust to three different parameter choices (**fig. S4**). To identify genomic features associated with boundary activity, we sorted boundaries by their insulation strength and searched for previously-mapped factors (*29*) that were correlated with boundary activity (**Figs. 3B-E**). Consistent with prior Hi-C analyses of domain folding (*6, 27*), we find architectural factors including RAD21, CTCF, YY1, and ZNF143 enriched at strong boundaries. More generally, boundaries are highly enriched for promoter marks and are localized precisely to nucleosome-depleted regulatory elements, consistent with the prior identification of TSSs as boundary elements in budding and fission yeast (*10, 11*). Boundary activity at CTCF binding sites and at promoters is also readily apparent as clearing of interactions in the upper-right/lower-left corners of the CTCF-and TSS-aligned heatmaps in **Fig. 2D**. Importantly, although binding of CTCF and YY1 are both strongly positively-correlated with boundary scores (**Fig. 3B**), these factors did not always co-occur at every boundary – **Fig. 3E** shows successive classification of boundaries into CTCF-associated boundaries, CTCF-negative YY1-enriched boundaries, CTCF-and YY1-depleted promoter boundaries, and a fourth class of weak boundaries largely depleted of all three features (**fig. S5**). Together, these data precisely localize chromatin domain boundaries in human cells, emphasizing the tight association between nucleosome depletion/dynamics (and/or the diverse set of proteins that occupy various nucleosome-depleted regions) as key players in insulating adjacent chromatin interaction domains from one another.

**Figure 3.**
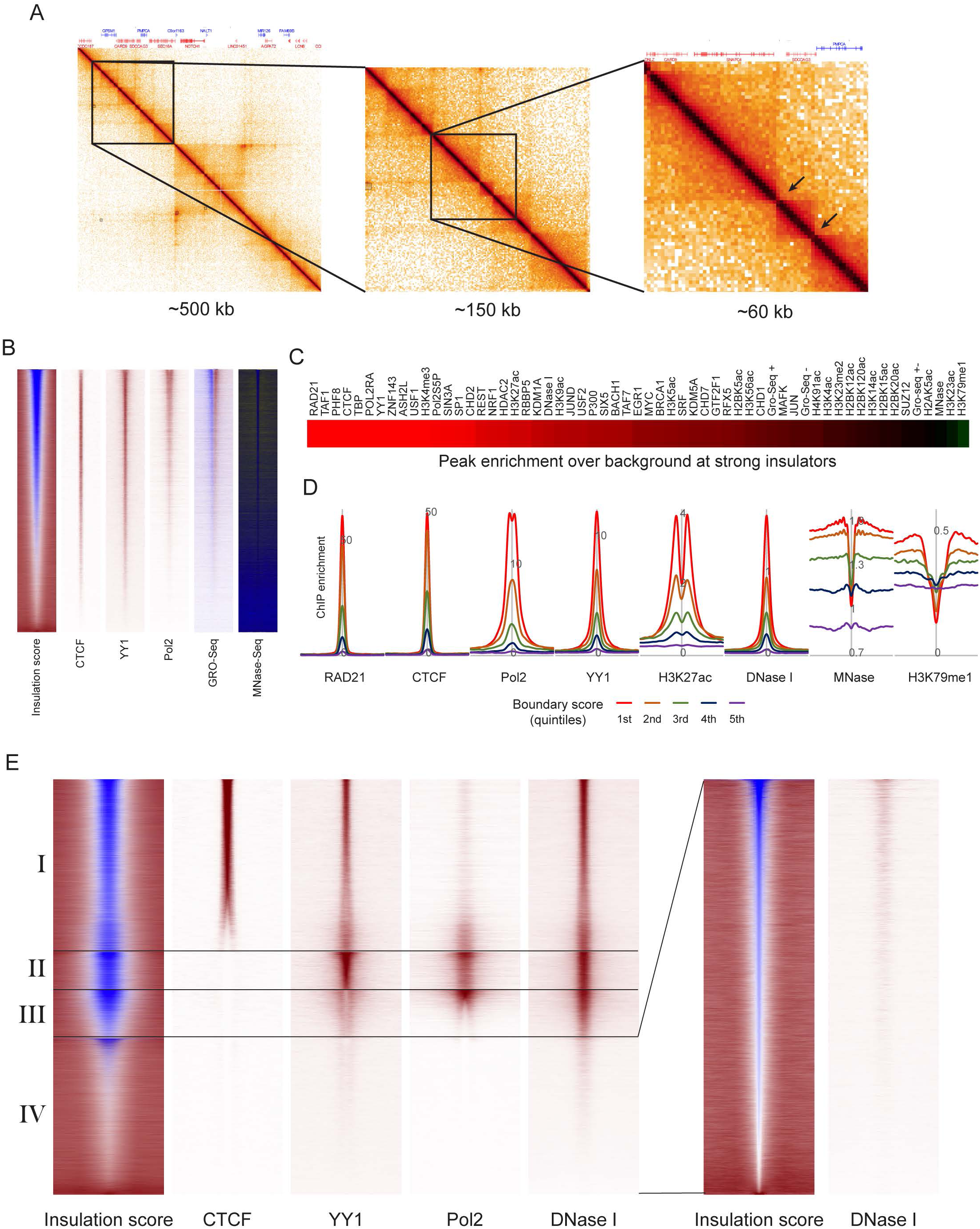
High resolution identification of interaction boundaries. (A) Successive zooms into the HFF Micro-C dataset show a boundary between self-associating domains, located at a promoter. (B) Gross features of boundary elements in ESCs (all systematic comparisons are performed in ESCs given the abundant ChIP-Seq data available in this cell type). Boundaries were identified as described in Methods, and the strongest 100,000 boundaries (**Methods**) are sorted according to boundary score (left panel). See also **fig. S4**. Right panels show ChIP-Seq enrichment for key boundary factors, DNase-and MNase-Seq data, and GRO-Seq as a readout of active transcription. (C) Survey of factors enriched at strong boundaries. For the indicated ChIP-Seq (and DNase-and MNase-Seq) datasets, the peak to trough (ChIP signal at the central 500 bp vs. the baseline, 2 kb distant) ratio was calculated, and factors are ordered by enrichment score. (D) Examples of boundary-enriched (CTCF, etc.) and -depleted (H3K79me2) factors. (E) Independence of key boundary elements. Boundaries were successively sorted by CTCF, YY1, and Pol2 to group boundaries into four classes – CTCF-associated, CTCF-negative/YY1-positive, CTCF/YY1-depleted/Pol2-positive, and weak CTCF/YY1/Pol2-depleted boundaries (see **fig. S5**).

Among other TAD-associated structural features, we noted that the most apparent difference between Hi-C and Micro-C maps is a profusion of dots in Micro-C that are indistinct or absent in Hi-C maps (**Figs.1A-B, Fig. 4A, fig. S6**). This intuition is confirmed computationally, as systematic identification of looping interaction peaks (**Methods**) reveals a massive increase in the number of dots and dot anchor loci in Micro-C maps relative to Hi-C (**Fig. 4B**). In general, the majority of dots detected in Hi-C libraries were also detected with Micro-C (86.6% in ESC, 88.9% in HFF), although even for these Hi-C/Micro-C common dots the Micro-C dataset exhibited increased signal-to-noise (**Fig. 4C, fig. S7**). Conversely, averaging interaction maps for Micro-C-specific dots revealed a low level of signal enrichment at these locations in the Hi-C dataset (**Fig. 4C, fig. S7**), indicating that evidence for these interaction peaks is present in both datasets but that Micro-C resolves more dots as a consequence of the improved signal-to-noise of this protocol.

**Figure 4.**
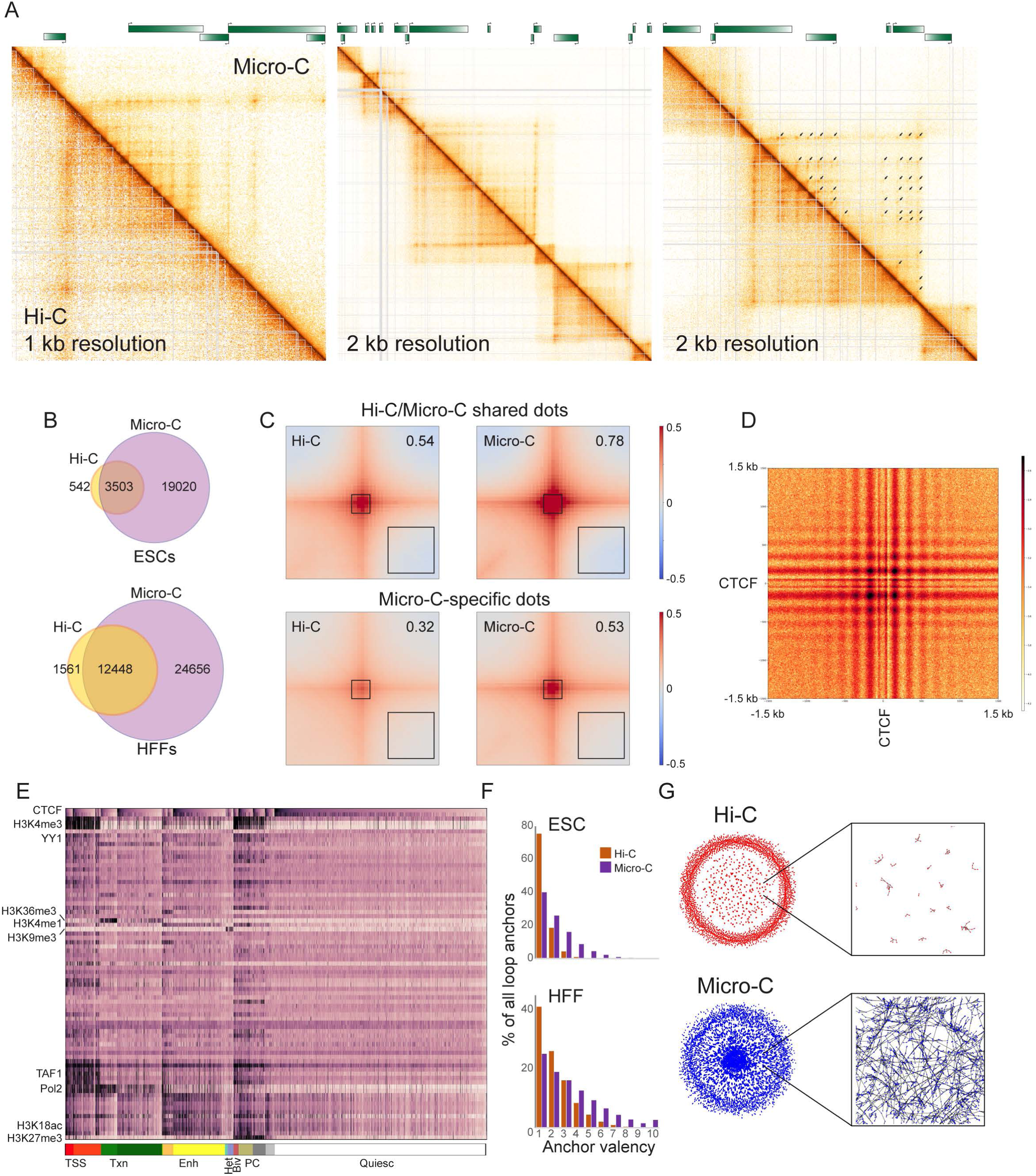
TADs are composed of a heterogeneous network of internal looping interactions. (A) Examples of Micro-C-specific peaks of looping interactions in HFFs. In each case Micro-C contact map is shown above the diagonal, along with corresponding Hi-C heatmap below the diagonal (see also **fig. S6** for ESC examples). (B) Venn diagrams show presumed looping interaction peaks (“dots”) identified by Micro-C and Hi-C in ESCs and HFFs, as indicated. (C) Heatmaps showing average contact frequency for dots called in both Hi-C and Micro-C (top), or in Micro-C only. Number in the upper right hand corner represents the signal strength at the loop base (heatmap center) above the nearby background (black box, lower right corner). See also **fig. S7** for HFF dataset. (D) Heatmap showing average contact frequency, at nucleotide resolution, for all pairs of CTCF ChIP-Seq peaks associated with convergently-oriented CTCF motifs at separations between 100 kb and 1Mb. (E) Global view of dot anchor sites. For all anchor sites, enrichment for various chromatin proteins or histone modifications (*29*) was computed. Anchor sites are first sorted according to chromatin state (broad chromatin types indicated at bottom, from left: Transcription Start Sites, Transcribed chromatin, Enhancers, Heterochromatin, Bivalent, Polycomb, and Quiescent), then sorted according to CTCF enrichment within each subcluster. See also **figs. S8-10**. (F) Micro-C identifies genomic loci with multiple looping interaction peaks. Histograms show the number of looping interaction peaks for any given genomic locus, revealing a clear shift towards multiple peaks in Micro-C compared to Hi-C datasets. See also **fig. S11**. (G) Grid completeness. Left panels show networks constructed from Hi-C (top) and Micro-C (bottom) looping interaction peaks, while right panels show a zoom in from the center of the network. Here, nodes represent genomic loci (anchors), while edges represent interaction peaks between anchor sites (dots).

Where are Micro-C-specific dots located? We noted that many examples of Micro-C dots were found along extrusion-associated stripes at TAD borders (**Fig. 4A**), suggesting the presence of multiple, relatively weak, pause sites that temporarily arrest SMC extrusion complexes before these complexes reach the “hard stop” boundaries previously observed at inwardly-oriented CTCF sites (*27, 30*). Moreover, we often observed Micro-C-specific dots populating the interior of TAD boxes, located at the intersection of the newly-identified weak pause sites found along TAD-bordering stripes. What is the nature of the newly-identified genomic dot anchors? Consistent with the central role for CTCF in blocking or pausing loop extrusion (*27, 30*), we find that the majority of new dot anchors coincide with ChIP-Seq peaks of CTCF enrichment, with∼77% of all dot anchors linked to CTCF binding sites. To explore ultrastructural features of CTCF-anchored looping interactions, we produced read-level pileup contact maps averaging off-diagonal pairs of convergently-oriented CTCF binding sites (**Fig. 4D**). Two salient features of these heatmaps are worth noting. First, the “beading” of the central interactions reveals interactions between the well-positioned nucleosomes flanking CTCF binding sites (**Fig. 2B**). Second, the horizontal and vertical lines on this heatmap are consistent with the stripes that characterize the loop extrusion process in Hi-C heatmaps. Importantly, the extension of these stripes beyond (above or to the right of) the anchor points is consistent with many of these CTCF-mediated dots occurring in the middle of a loop extrusion track, generalizing our observation that many Micro-C-specific dots are identified along these extrusion stripes (see, e.g, **Figs. 1A** and **4A**).

To explore the molecular features of these enriched looping interactions, we calculated the enrichment of a variety of structural proteins and histone modifications at dot anchors, then clustered dot anchors according to these features (**Fig. 4E, fig. S8**). We find that multiple classes of genomic loci are involved in looping interaction peaks, including: 1) Promoters (enrichment for H3K4me3, Pol2); 2) Enhancers (H3K4me1); 3) Coding regions (H3K36me3, Pol2); 4) Polycomb chromatin (H3K27me3, SUZ12); 5) “Quiescent” chromatin (depleted of distinguishing histone marks). Beyond the expected enrichment of CTCF at dot anchors, we identify a number of CTCF-depleted dot anchors, although interestingly these anchors were generally found at the same types of genomic element as the CTCF-enriched anchors (**fig. S9**). Given that systematic identification of dot anchors is somewhat inefficient – for instance, Micro-C-specific dots clearly exhibit signal in Hi-C but are not called – we further explored CTCF-independent dots by averaging interactions between various pairs of CTCF-depleted genomic loci (**fig. S10**). These heatmaps reveal strong Micro-C signal enrichment for enhancer-promoter interactions (**fig. S10A**), as well as dots occurring between paired binding sites for a wide array of transcriptional and chromatin regulatory proteins (**fig. S10B**).

Finally, we noted that most of the new dots identified by Micro-C were relatively weak interaction peaks occurring either at the intersection points of new anchor sites in the interior of TADs, or along stripes associated with strong CTCF dots at TAD corners. This suggests that a given genomic locus can participate in multiple distinct transient loops across a cell population. To more systematically evaluate these qualitative observations, we first plotted the number of dot interactions for each genomic location involved in at least one dot. Consistent with visual examination, we find a significant increase in the fraction of dot anchors that participate in more than one interaction in Micro-C, relative to Hi-C (**Fig. 4F, fig. S11**). The implication of this finding is that Micro-C reveals a more extensively interconnected network among genomic loci. Indeed, visualization of connectivity networks derived from Hi-C and Micro-C data (**Fig. 4G**) revealed a more densely connected structure for the Micro-C dataset. In graph theory terms, this can be quantitated as node transitivity (*31*), with the Micro-C network exhibiting significantly greater transitivity than the Hi-C network (0.46 vs. 0.29 for ESCs, 0.57 vs 0.48 for HFFs), confirming that Micro-C dot interactions form a more complete grid than do Hi-C dots.

Taken together, our data reveal that the Micro-C protocol yields improved signal-to-noise, as well as superior genomic resolution, relative to Hi-C. Biologically, our data extend our understanding of chromosome folding in mammals in several ways. We localize boundaries between contact domains precisely to nucleosome-depleted regions, consistent with the location of contact domain boundaries in budding yeast (*10*), and suggesting that multiple distinct factors can interfere with interactions between adjacent chromatin domains. Our data also dramatically increase the list of genomic loci involved in presumed looping interactions, updating our view of TADs as heterogeneous populations made up of transient loops formed by multiple weak pause sites that slow or stall SMC protein movement. This internal structure is “blurred out” by noise and the coarser capture radius of restriction enzyme Hi-C, but is uniquely resolvable by Micro-C.

## ACKNOWLEDGEMENTS

We thank A. Goloborodko for critical discussions. All authors acknowledge support from the National Institutes of Health Common Fund 4D Nucleome Program (U54-DK107980). NK is supported by HFSP grant LT000631/2017-L.

## SUPPLEMENTARY MATERIALS

Materials and Methods

Figs. S1 to S11

Tables S1 to S4

