## Supplement for "Ultrastructural details of mammalian chromosome architecture"

### Materials and Methods

#### Cell culture

Human Embryonic stem cells (H1 – WiCell, WA01, lot # WB35186) were cultured in mTeSR1 media (StemCell Technologies, 85850) under feeder-free conditions on Matrigel hESC-qualified matrix (Corning, 354277, lot # 6011002) coated plates at 37$^{o}$C and 5% $CO_2$. H1 cells daily fed with fresh mTeSR1 media and passaged every 4-5 days using ReLeSR (StemCell Technologies, 05872) reagent. Cells were dissociated in to single cells with TrypLE Express (Thermo Fischer Scientific, 12604013) for the downstream applications.

#### Generation of *in-situ* Hi-C libraries with DpnII

Deep Hi-C libraries were prepared following the Hi-C 2.0 protocol as previously described (*1*). Briefly, for each deep library, 4x 5 million cells were crosslinked with 1% formaldehyde for 10 minutes at room temperature. Cells were lysed and chromatin was digested overnight at 37°C with DpnII (NEB), In situ ligation of was performed for 4 hours at 16°C with T4 DNA ligase (Invitrogen) on ends that were blunted in the presence of biotin-14-dATP (Invitrogen). DNA was isolated after reverse crosslinking at 65°C with proteinase K (Sigma). Then, biotin was removed from unligated ends at 23°C using Klenow enzyme (Invitrogen) in the absence of dTTP and dCTP. DNA was fragmented to 200-300bp by Covaris sonication followed by a double size selection with SPRI beads (AMpure XP, Beckman Coulter) to select for DNA of 250-350bp. After end repair, biotin labeled DNA was pulled down by streptavidin coated MyOne C1 beads (Invitrogen) on a magnetic stand. Illumina paired-end sequencing adaptors were ligated to both ends at room temperature for 2 hours (T4 DNA ligase, Invitrogen) and PCR was performed using Q5 polymerase (NEB) to generate the final Hi-C library. Primers were removed with SPRI beads and 4 tubes were pooled prior to sequencing. Typically, deep libraries were generated from a minimum of 8 lanes of sequencing using our in-house Illumina Hi-seq 4000.

#### MicroC-XL

The Micro-C XL protocol was adopted from (*2*). Frozen cells were resupended in 200 ul cold 1x PBS (10 nM $Na_2HPO_4$, $KH_2PO_4$, pH 7.4, 137 nM NaCl, 2.7 nM KCl,) per 1 mio cells and split into 1 mio cells aliquots. Note, 1x BSA (NEB, #B9000S) was added to PBS prior resuspension and wash to reduce stickiness of HFF cells to the tub walls. After 20 min incubation on ice, cells were collected by centrifugation (5000x g, 5 min), washed with 500 µl buffer MB#1 (10 mM Tris-HCl, pH 7.5, 50 nM NaCl, 5 mM MgCl2, 1 mM $CaCl_2$, 0.2% NP-40, 1x Roche cOmplete EDTA-free (Roche diagnostics, 04693132001)), collected by centrifugation (5000 g, 5 min), and resuspended in 200 µl MB#1. Chromatin was incubated with MNase for 20 min at 37°C. MNase concentrations were chosen to yield mostly mono-nucleosomal fragments, as tested in prior digestion tests, typically 5-20 U MNase (Wortington Biochem, LS004798). The digestion was stopped by addition of 0.5 mM EGTA (Bioworld, #405200081) to a 1.5 mM final concentration and incubation at 65°C for 10 min.

Chromatin aliquots were pooled for further processing. Here, the equivalent of 2.5 mio cells input yielded the best results, more than 5 mio cell-equivalent per aliquot is not recommended. The chromatin was collected by centrifugation (5000x g, 5 min), washed with 500 µl 1x NEBuffer 2.1 (NEB, #B7202S), collected by centrifugation (5000x g, 5 min), and resuspended in 45 µl NEBuffer 2.1. DNA ends were dephosphorylated by addition of 5 µl rSAP (NEB, #M0203) and incubation at 37°C for 45 min. The reaction was stopped by incubation at 65°C for 5 min. 5' overhangs were generated by 3' resection. Here, 40 µl pre-mix (5 µl 10x NEBuffer 2.1, 2 µl 100 mM ATP (Thermo Fisher, #R0441), 3 µl 100 mM DTT, 30 µl H2O) and 8 µl Large Klenow Fragment (NEB, #M0210L) and 2 µl T4 PNK (NEB, #M0201L) were added to the sample in respective order. The reaction was incubated at 37°C for 15 min. The DNA overhangs were filled with biotinylated nucleotides by addition of 100 µl pre-mix (25 µl 0.4 mM Biotin-dATP (Invitrogen, #19524016), 25 µl 0.4 mM Biotin-dCTP (Invitrogen, #19518018), 2 µl 10 mM dTTP and 10 mM dTTP (stock solutions: NEB, #N0446), 10 µl 10x T4 DNA Ligase Reaction Buffer (NEB #B0202S), 0.5 µl 200x BSA (NEB, #B9000S), 38.5 µl H2O) and incubation at 25°C for 45 min. The reaction was stopped by addition of 12 µl 0.5 M EDTA (Invitrogen, #15575038) and incubation at 65°C for 20 min.

The chromatin was collected by centrifugation (10000x g), washed in 500 µl 1x Ligase Reaction Buffer, and collected by centrifugation (10000x g). The chromatin pellet was resuspended in 2500 µl ligation reaction buffer (1x NEB Ligase buffer, 1x NEB BSA, 12500 U NEB T4 Ligase (NEB, #M0202L)) and incubated rotating at RT for 2.5-3 h. After proximity ligation, the chromatin was collected and resuspended in 200 µl 1x NEBuffer 1 (NEB, #B7001S) and 200 U NEB Exonuclease III (NEB, #M0206S). For deproteination and reverse crosslinking, 25 µl ProteinaseK (25mg/ml in TE with 50% glycerol) and 25 µl 10% SDS (Invitrogen, #15553-035) were added and the sample was incubated at 65°C o/n.

The DNA was first phenol/chloroform purified and second purified with DNA Clean & Concentrator Kit (Zymo, #D4013). The 300 bp sized MicroC library was purified via 1.5 % agarose gel electrophoresis and extracted with Zymoclean Gel DNA Recovery Kit (Zymo Research, #D4002) with a final elution volume of 50 µl.

5 µl DynabeadsTM MyOneTM Streptavidin C1 beads (Invitrogen, #65001) were washed twice with 300 µl 1x TBW (5 mM Tris-HCl, pH 7.5, 0.5 mM EDTA, 1 M NaCl) and suspended in 150 µl 2x TBW (10 mM Tris-HCl, pH 7.5, 1 mM EDTA, 2 M NaCl). 100 µl $H_2O$ and 150 µl washed Streptavidin beads in 2x TBW were added to the sample and incubated rotation at RT for 20 min. The beads were washed twice with 300 µl 1x TBW and resuspended in 50 µl TE buffer. Sequencing libraries were prepared with NEBNext® Ultra™ II DNA Library Prep Kit for Illumina® (NEB, #E7645) according to protocol, except for the DNA purification and size selection prior PCR. Here, adaptor-ligated DNA was pulled-down via bound Streptavidin beads and washed twice with 300 µl 1x TBW and once with 0.1x TE. Finally, the beads were resuspended in 20 µl 0.1x TE. PCR amplification, sample indexing, and DNA purification after PCR was performed with according to (NEB, #E7645) using NEBNext® Multiplex Oligos for Illumina®. The samples were sequenced on an Illumina HiSeq 4000 on 50 base pair paired end mode.

### Data analysis

#### Chromosome Conformation Capture data processing

Hi-C and Micro-C datasets were processed with the *distiller* pipeline (https://github.com/mirnylab/distiller-nf). Briefly, reads were mapped to the human reference assembly hg38 using bwa mem with flags -SP. Alignments were parsed and pairs were classified using the pairtools package (https://github.com/mirnylab/pairtools) to generate 4DN-compliant pairs files. Pairs having matching alignment strands and coordinates with a possible 1 bp offset were considered PCR and/or optical duplicates and thus removed.

Pairs classified as uniquely mapped or rescued chimeras with high mapping quality scores on both sides (MAPQ > 30) were aggregated into contact matrices in the cooler format using the *cooler* package (https://www.biorxiv.org/content/10.1101/557660v1) at 500bp and into multiresolution cooler files (500bp, 1kb, 2kb, 5kb, 10kb, 50kb, 100kb, 250kb, 500kb, 1Mb). All contact matrices were normalized using the iterative correction procedure (*3*) after bin-level filtering. Low-coverage bins were excluded using the MADmax (maximum allowed median absolute deviation) filter on genomic coverage, described in (*4*), using a threshold of 5.0 MADs. To remove short-range ligation artifacts at a given resolution—such as unligated and self-ligated molecules—we removed pairs mapping to the same or adjacent genomic bins.

**Contact heatmaps**

The dense matrix data was extracted from balanced cooler files with *cooltools* package in python and plotted with *grid* package in R. The maximum color intensity was set to the percentile that allowed the best assessment of matrix specific features, such as TADs or dots. For heatmaps that display MicroC and HiC within one heatmap, the value-to-color conversion was applied to the individual data sets for equal percentiles.

**Eigenvectors**

Compartmentalization was assessed using an eigenvector decomposition procedure based on (*3*), as implemented in the *cooltools* package (https://github.com/mirnylab/cooltools). Eigenvector decomposition was performed on observed-over-expected *cis* contact maps at 100-kb resolution, separately for each chromosomal arm, after subtracting 1. The first three eigenvectors and eigenvalues were calculated, and the eigenvector associated with the largest absolute eigenvalue was chosen. We used an identically binned track of gene frequency to orient the eigenvectors such that the positive and negative values of the eigenvector correlate with high and low gene density, respectively.

**Interaction decay**

Scaling curves of contact frequency as a function of genomic separation were generated by aggregating normalized contact frequency over valid pixels along diagonals of 1kb-resolution cis contact maps, with diagonals grouped into geometrically increasing spans of genomic separation. Scaling curves were also generated for unbinned data (uniquely mapped paired-end reads) using a similar methodology, independently for reads having each of four possible strand orientations. Average contact frequency P(s) curves are displayed using log-log axes.

Scaling curves of contact frequency as a function of genomic separation for short distances were generated from valid pairs files. Here, the distance between the read starts of uniquely mapped, cis reads were extracted and normalized for a distance of 1 to 100 kb to 1. For read shifted interaction decay curves, the read start were shifted by 73 bp with respect to read orientation.

### Insulation score

Diamond insulation scores (*5*) were calculated on 1-kb resolution maps using the *cooltools* package. A local minima detection procedure based on peak prominence, described in (*6*) and implemented in *cooltools*, was used to call insulating loci.

The diamond insulation (*5*) was computed for 100 and 200 bp resolutions. Here, 100 bp and 200 bp binned, balanced multi-cooler files were generated form valid pairs files with reads shifted by 73 bp to the theoretical nucleosome dyad. The insulation score was computed with the *cooltools insulation* tool at 100 bp resolution with 1,000 and 10,000 bp diamonds and for a 200 bp resolution with a 2,000 bp diamond. The positions for the strongest 100,000 boundaries was extracted as alignment point for 1D analyses.

### 1D analysis

Genomic alignment points: The positions for the strongest 100,000 boundaries, transcription start site, or CTCF peak positions, were used as alignment points for 1D analysis.

Signal tracks: For nucleosome occupancy data, the first read of the Micro-C read pairs was mapped with Bowtie2 (2.3.2) and Samtools (1.4.1) against the human genome (hg38) using the following parameters (bowtie2 -p 16 --local -x {hg38 bowtie2 genome} -U {fastq.file} | samtools view -bS -o {output.bam} -) 2>&1 | tee {report.file}). To compute the nucleosome dyad density, the resulting read starts were shifted by 73 bp (strand sensitive) to obtain the theoretical nucleosome dyad. ChIP-seq signal data tracts were downloaded from ENCODE (see **Table S2**). Raw Gro-Seq data was downloaded from and processed as described in (*7*).

Sorting of 1D heatmaps and computation of quintals: The mean PolII ChIP-seq signal in a window form –150 to 250 bp around TSSs was computed with respect to the direction of transcription, was used as a proxy for transcription rate. For CTCF binding strength, the mean CTCF ChIP-seq signal around CTCF peaks (ENCFF368LWM) was computed in a window of –100 to 100 bp. Similarly, the mean insulation score within a window of –100 to 100 bp at boundary sites was used to sort alignments at boundaries. Alignment sites with NA scores within a window of -100 to 100 bp around the alignment point were removed from analysis, as they represent unmappable regions within the genome. Quintiles were computed by extracting 5 equally sized lists of alignment points from sorted data.

Plotting of 1D heatmaps and average signal plots: The signal around alignment points for given samples were extracted within a -2000 to 2000 bp window. The extracted data was mean-binned for 25 bp bins and visualized with gridGraphics package in R. For average signal plots, the data was prepared as for heatmaps, but the mean signal for 25 bp bins was computed for given quantiles and plotted.

### Contact frequency pileup maps

We used *cooltools* to calculate aggregate contact frequency maps (pileup maps) centered at specific genomic loci. For lists of single or paired genomic landmarks, pileup maps consist of aggregated contact map “snippets” encompassing fixed-length genomic windows along each axis, centered at the midpoints of the respective features. These snippets are extracted from either iteratively-corrected or observed-over-expected contact matrices or from unbinned contacts. For unbinned data, the coordinates of the 5’ ends of the alignments from each uniquely mapped molecule were shifted 73 bp along the alignment reference strand (i.e. to the approximate locations of the nucleosome dyad axes) before aggregating.

Due to the localization uncertainty associated with dot calls, which were performed on 5- and 10-kb contact maps, the pileup maps for those features were generated using 5-kb contact maps. For more precise genomic landmarks, pileup maps were generated using 500bp resolution contact maps. The pileup maps generated from unbinned data (paired-end reads) were centered on motifs associated with CTCF binding sites. This list of sites was created by intersecting CTCF ChIP-seq peaks from ENCODE (ENCFF368LWM) with a list of CTCF motifs (JASPAR motif ID #MA0139.1) using *bedtools* (*8*).

**Dot detection**

We reimplemented the HiCCUPS algorithm (*9*) in the form of the *call-dots* command line tool as a part of the "cooltools" package of tools for Hi-C data analysis (https://github.com/mirnylab/cooltools). This tool was used to identify locally enriched interactions, i.e. "dots", both for Hi-C and Micro-C binned data. Dots were called at 5kb and 10kb resolutions using only high quality mapped pairs (MAPQ > 30) and the resulting lists were merged using the same criteria as in (*9*):

- matching dot calls were identified between 5kb and 10kb lists, assuming dots with Euclidian distance between them lower than 10kb as matching
- all unique dots called at 10kb were retained in the merged list
- only higher resolution, 5kb, dot-calls were retained, for the matching 5kb and 10kb calls
- unique dot-calls at 5kb were retained only in case they were small (<100kb) in size, or particularly strong (>=100 raw interactions per 5kb pixel)

A detailed description of the dot calling procedure for a given resolution is provided in (*9*); here we briefly highlight some of the major steps and the deviations from the original procedure (*9*) that we assumed here:

- detecting locally enriched interactions, i.e. dot-calling, starts with defining locally-adjusted expected level of interactions for pixels on a heatmap
- in this case, we limit ourselves to survey only pixels that are separated by less than 10Mb, i.e. 10Mb-wide band near the diagonal on the heatmap was surveyed.
- for a given pixel, its locally-adjusted expected defined as the average distance decay corrected by the enrichment of the area surrounding the pixel of interest, i.e. P(s)*donut_observed/donut_expected
- like in HiCCUPS, we employ 4 independent filter-shapes: "donut", low-left, vertical, and horizontal
- our calculation of local enrichment factor deviates from the original HiCCUPS - we do not adjust the footprint of the filters neither for pixels near the diagonal nor for pixels in sparse areas of the heatmap, instead we convolve a fixed size filters/kernels with the observed/expected heatmaps to calculate local enrichment factors
- surveyed pixels are grouped into so-called lambda-chunks according to their level of locally-adjusted expected and BH-FDR is performed for each of lambda-chunks independently, as described in (*9*).
- pixels identified as significant in each lambda-chunk constitute the filtered list of peak calls that gets post-processed the same way as in the original HiCCUPS procedure
- clustering is done using Birch initialization procedure and additional enrichment thresholding is performed along with the special treatment of the non-clustered singletons.

### Dot anchor analysis

Dot calls between Hi-C and Micro-C were compared using the following methodology. First, for each individual list of dot calls (Hi-C and Micro-C), every call was assigned a unique protocol-specific dot-call ID. Then, the left (5'-most) anchor intervals from the two call lists were intersected using *bedtools intersect*, such that every resulting intersection was associated with a pair of dot-call IDs (one from each protocol). The same procedure was applied to intersect the right (3'-most) anchor intervals from the two call lists. Common dots were identified from matching pairs of dot-call IDs found in the two intersection sets. The remaining dot calls were identified as unique to their respective protocol of origin. To account for uncertainty in dot localization, we increased the genomic extent of each dot anchor interval by 10kb in each direction before performing the intersections.

Distinct anchor regions from dot calls were assessed using a similar methodology. First, for each protocol, overlapping anchor intervals across all dot calls were grouped using *bedtools cluster* with -d 20000 and those groups were merged to produce a list of non-overlapping consensus intervals. The consensus anchor interval lists from Hi-C and Micro-C were then combined and the intervals grouped using the same command. The groups were then merged to produce a final consolidated list of non-overlapping anchor intervals. Those final anchor intervals derived from only one of the two protocols were identified as being unique to that protocol, while the remaining consolidated intervals were identified as being common to both protocols.

### Dot anchor graph analysis

Networks were constructed using lists of dots, where dot anchors represent nodes while dot calls represent edges between nodes. Such a representation allows us to leverage the properties of networks for the comparison of the lists obtained from the two protocols. Specifically, we look at the degree distribution and clustering coefficient of the network.

The degree distribution of the network represents the distribution anchor valencies. The network is constructed, and the degree of each node is extracted. A histogram of the degrees can be generated for each network and compared.

The clustering coefficient is a frequently used property of networks that measures the average interconnected-ness of the local neighborhood of around nodes. The clustering coefficient is calculated as the 3 times the ratio of closed triangles to open paths of length 2 found in the network. A fully connected graph would have a coefficient of 1. For the network of dots calls, distance dependence of dots ensures that its network will never be fully connected. To account for this, we adjust the transitivity by dividing it by the transitivity of a network that is fully connected up to 2Mb generated from all the anchors associated with the dots.

### CTCF occupancy of dot anchors

Dot anchors were intersected with CTCF ChIP-seq peaks using *bedtools* intersect. Since the genomic regions associated with dot anchors were extended during the clustering and accumulation process, no additional margins were introduced when intersecting with CTCF peaks.

For both classes of dot anchors (with a CTCF peak and without a CTCF peak), the mean fold change of CTCF ChIP-seq over the anchor region was also calculated and recorded and histograms were constructed.

### Epigenetic characterization of ESC dot anchors

Each distinct dot anchor interval identified in H1 hESCs was classified by its modal ChromHMM state assignment from the Epigenomics Roadmap 18-state model (trained on six core histone marks, 200bp bins) (*10*). Anchor intervals were further characterized by calculating the mean ChIP-seq signal in 10 kb windows around their midpoint for 79 histone and transcription factor experiments on H1 cells obtained from the ENCODE data portal. These feature vectors were concatenated into a heatmap, where each row corresponds to a dot anchor. Columns of the heatmap were sorted based on Ward agglomerative clustering (n=6) and rows were grouped by ChromHMM state. Clustering the rows using K-means or agglomerative clustering produced similar epigenetic classifications of the dot anchors.

**Looping interactions at enhancer-promoter and CTCF-depleted loci**

Enhancer locations for hg19 were downloaded from (http://zlab-annotations.umassmed.edu/enhancers/) and converted to hg38 with UCSC lift over tool (https://genome.ucsc.edu/cgi-bin/hgLiftOver). Intersections were computed with with bedtools window –w 100000 function and intersections with distances smaller than 5 kb were removed.

Processed ChIP-seq peaks for chromatin binding factors were extracted from ENCODE (**Table S3**). ChIP-seq peaks that are within 10 kb of a CTCF peak were removed with bedtools window –w 10000 –v function. Intersections of ChIP seq peaks within a ChIP seq data set were generated with bedtools window –w 100000 function. Sites where the two intersection resulting anchors were closer than 5 kb were removed.

As above, aggregate contacts were extracted with cooltools snippets function observed-over-expected contact matrices.

### SUPPLEMENTARY FIGURES

**Fig. S1. Micro-C and Hi-C maps of human pluripotent and differentiated cell types**

(A) Interaction decay curves for Micro-C maps of ESCs. Here, read pairs are separated according to their relative orientation. Note the excess of IN-IN interactions at short distance, representing undigested dinucleosomes.

(B) Zoom in of panel (A).

(C) As in panel (B), but with reads shifted 73 bp to the nucleosome dyad, thereby aligning IN-IN/IN-OUT/OUT-OUT read pairs.

(D) Step-wise decrease in interactions between adjacent nucleosomes. Each bar shows the ratio of ligation product abundance across two adjacent nucleosomes – the first set of bars shows the ratio of N/N+1 over N/N+2. Bars are shown for IN/IN, IN-OUT, OUT-IN, and OUT-OUT orientations. Notably, there is a greater dropoff from N+1 to N+2, and from N+3 to N+4, than there is between the nucleosome pairs N+2/3 and N+4/5, consistent with a zig-zag fiber architecture. These data suggest that in humans, in contrast to budding yeast, compacted chromatin fiber may extend beyond tetranucleosomes to organize longer stretches of the genome. Nonetheless, there is a marked dropoff from N/N+2 to N/N+4 and again from N/N+4 to N/N+6 – whether this represents inefficient extension of chromatin fiber compaction beyond the tetranucleosome, or a technical inability to recover N/N+4 products in a zig-zag fiber thanks to the interposed N+2 nucleosome, remains to be determined, although several independent analyses of in vivo chromatin folding (*11, 12*) support the former view: that chromatin forms fairly short, heterogeneous “clumps” of zig-zag fiber in vivo.

Figure S1

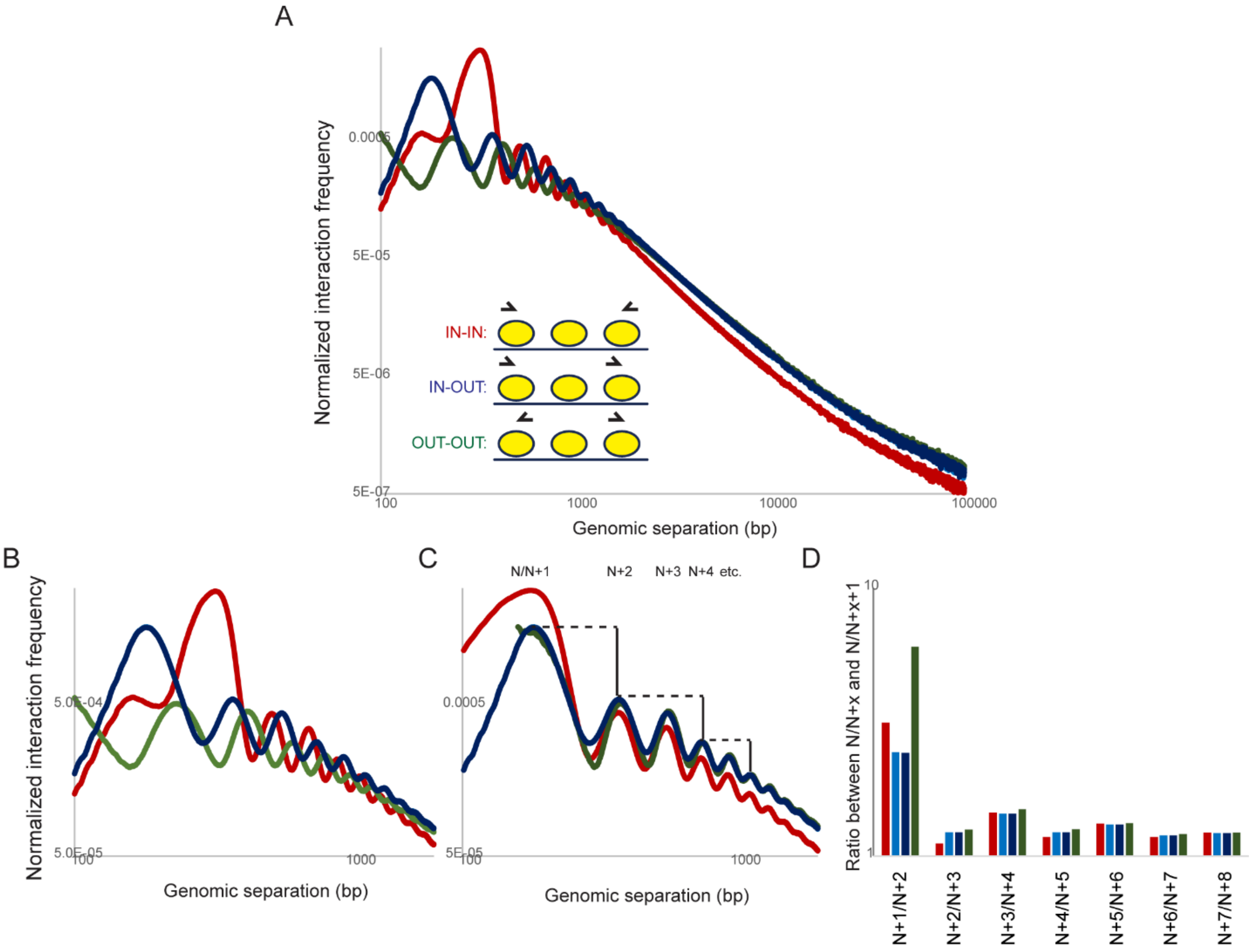
A
Normalized interaction frequency
0.0005
5E-05
5E-06
5E-07
100
1000
10000
100000
Genomic separation (bp)
IN-IN:
IN-OUT:
OUT-OUT:
B
Normalized interaction frequency
5.0E-04
5.0E-05
100
1000
Genomic separation (bp)
C
N/N+1
N+2
N+3 N+4 etc.
0.0005
5E-05
100
1000
Genomic separation (bp)
D
Ratio between N/N+x and N/N+x+1
10
1
N+1/N+2
N+2/N+3
N+3/N+4
N+4/N+5
N+5/N+6
N+6/N+7
N+7/N+8

**Fig. S2. Distribution of restriction sites influences Hi-C coverage**
Plots show frequency of Hi-C or Micro-C interactions > 10 kb (y axis) for genomic 2 kb bins with varying numbers of DpnII target sequences (x axis), revealing strong bias for poor Hi-C coverage for genomic intervals depleted of DpnII target sites.

Figure S2

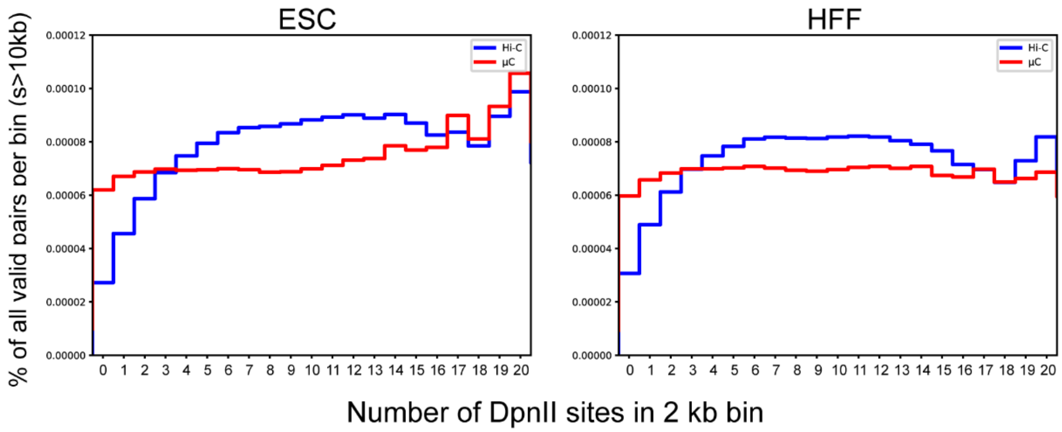

**Fig. S3. Increased compartmental organization of HFFs**

(A) Single-gene scale compartment. Left panel shows a broad zoom (chr11: 27,384,310-33,423,664) showing compartment signature (enriched interactions at long distance with the compartment checkerboard pattern) for this gene, while right panel shows the gene-scale zoom-in.

(B) Micro-C contact maps for ESCs and HFFs (above and below the diagonal, respectively) for the indicated chromosomes. The characteristic plaid compartment pattern is clearly stronger in HFFs compared to ESCs.

Figure S3

A

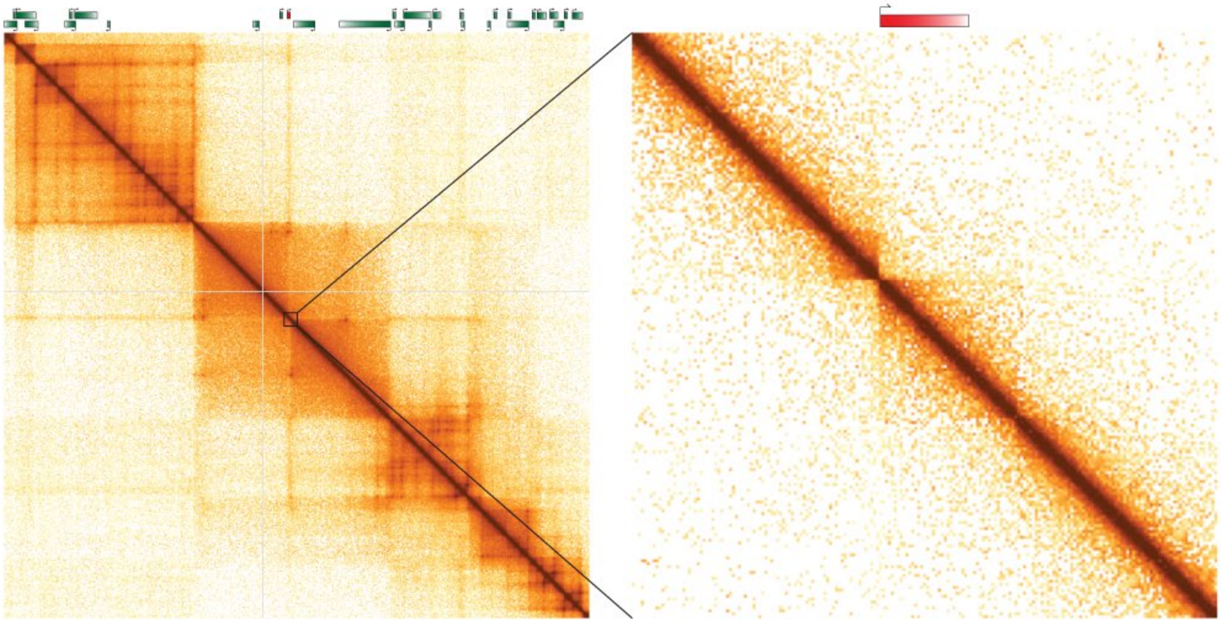

B

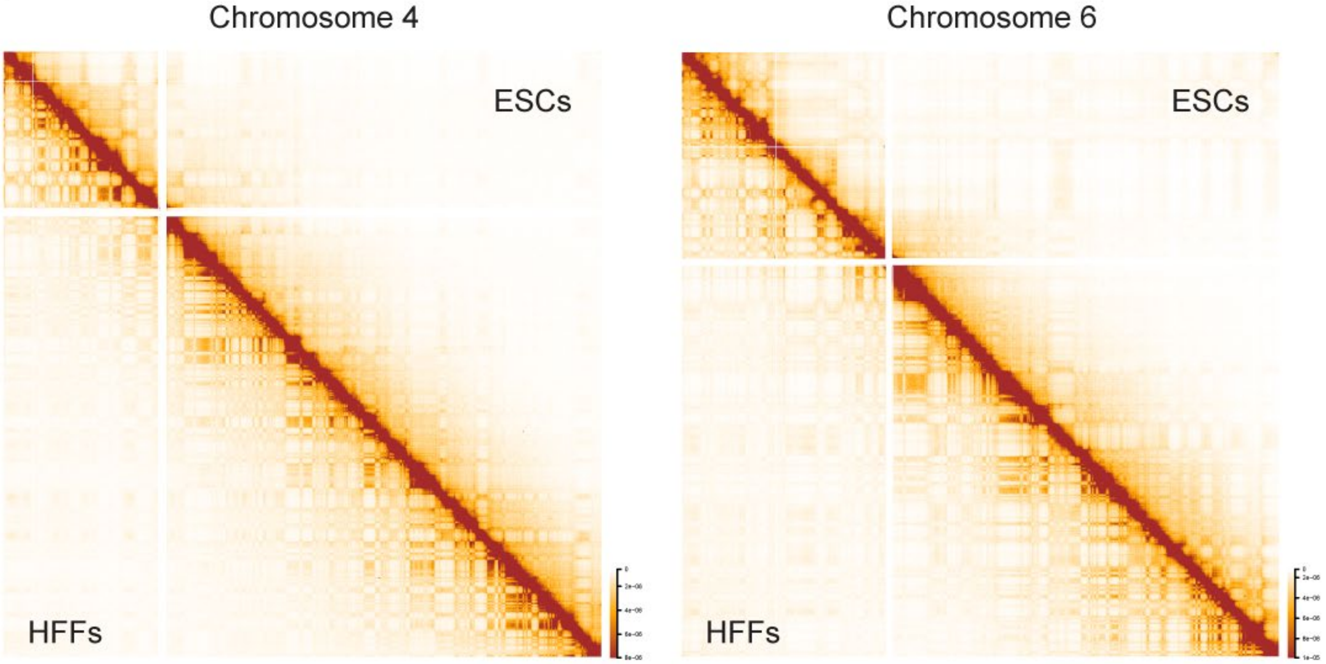

**Fig. S4. Boundary calls are not sensitive to parameter choice**
Boundaries were identified using a sliding window approach in which, for any given locus, the number of crossing interactions was calculated within some distance, relative to interaction frequency in windows on either side of the locus. Boundaries are identified as local minima in this vector. This approach has two free parameters: data bin size, and width of the sliding window. Here, boundaries were calculated using one set of parameters (100 bp bins, 1000 bp window), and boundary scores are shown in heatmaps for these and two additional parameter choices.

Figure S4

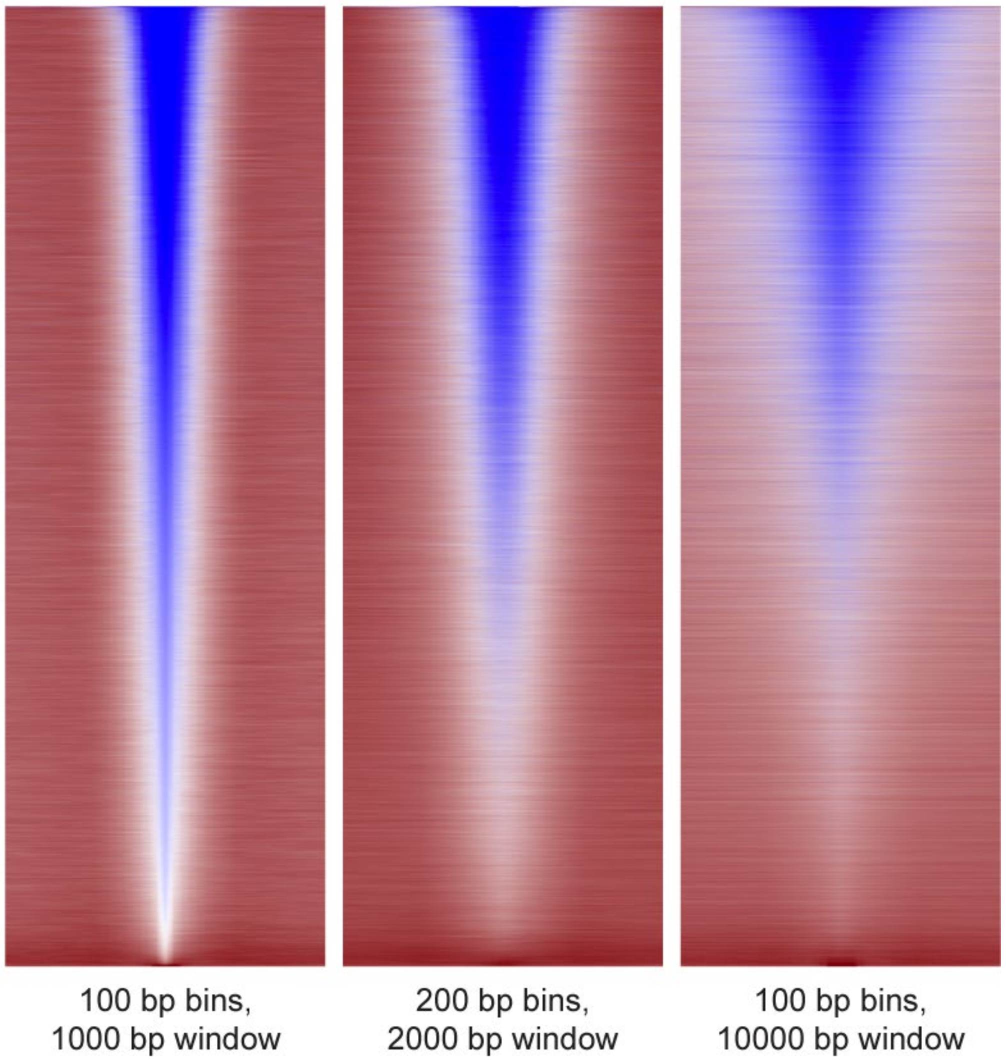

**Fig. S5. Molecular features of weak boundaries**
Heatmaps show ChIP (or DNase) signal for category IV boundaries from **Fig. 3E**. Several features, notably nuclease hypersensitivity, are associated with weak boundary elements.

Figure S5

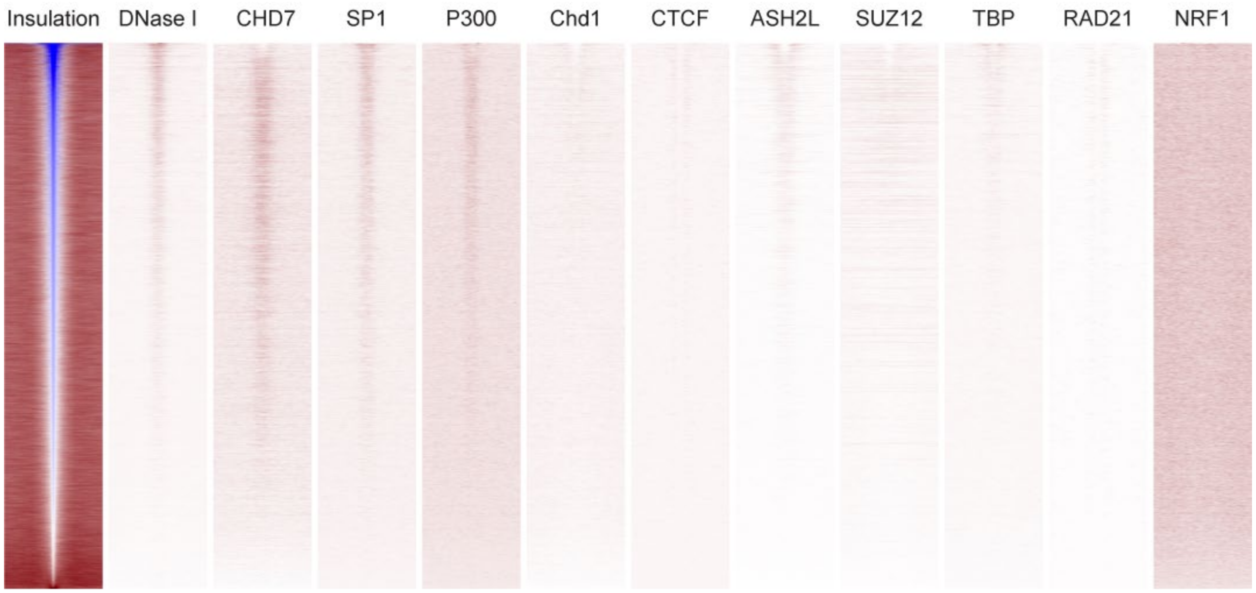

**Fig. S6. Additional examples of Micro-C-specific looping interactions**
Heatmaps show two examples of broad genomic regions (left: Chr8 98,000,000-100,200,000; right: chr6 127,979,027-131,127,602) exhibiting a number of Micro-C-specific loops in ESCs. Data tracks underneath show ChIP-Seq enrichment for key architectural and regulatory factors.

Figure S6

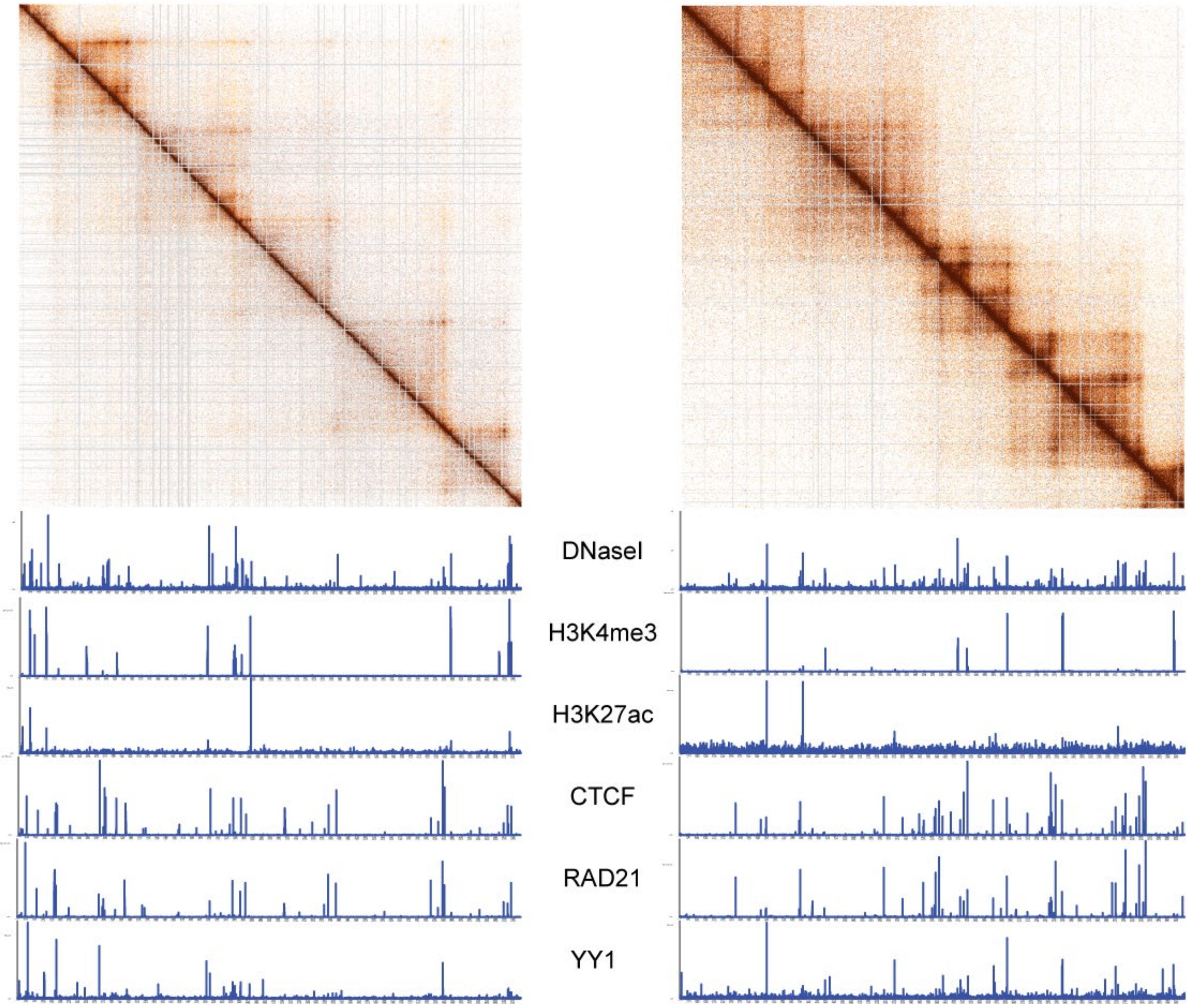

**Fig. S7. Loop averages for HFFs**
Data are shown as in **Fig. 4C** for the HFF dataset.

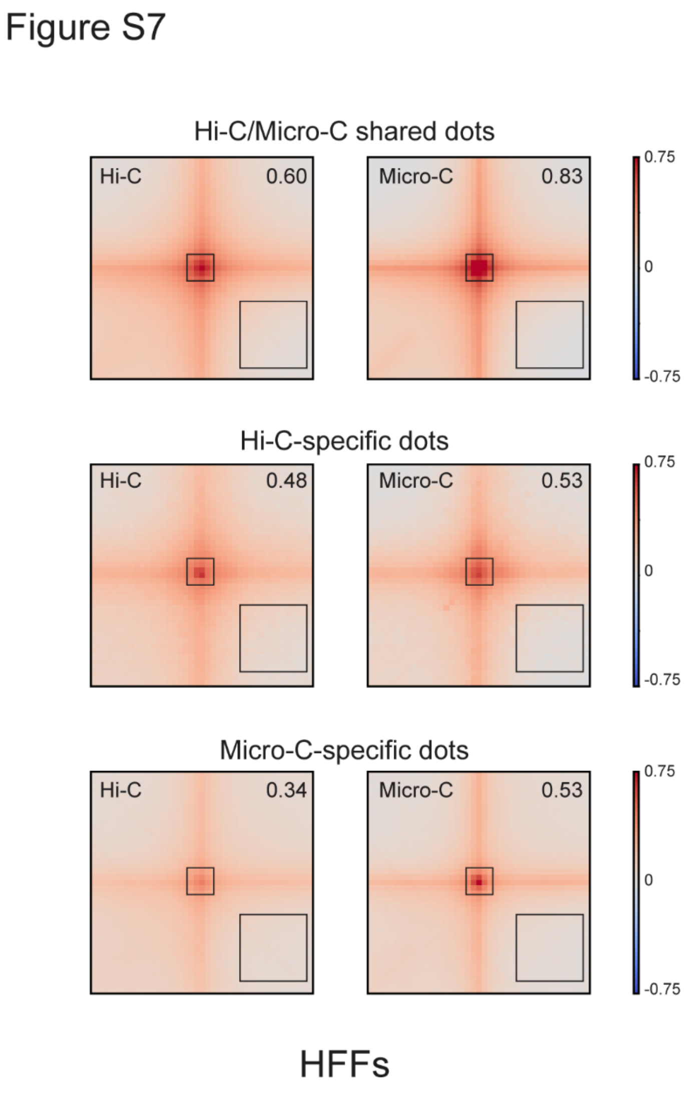

**Fig. S8. Molecular characteristics of loop anchors**
Heatmap from **Fig. 4E** is reproduced here in greater detail

Figure S8

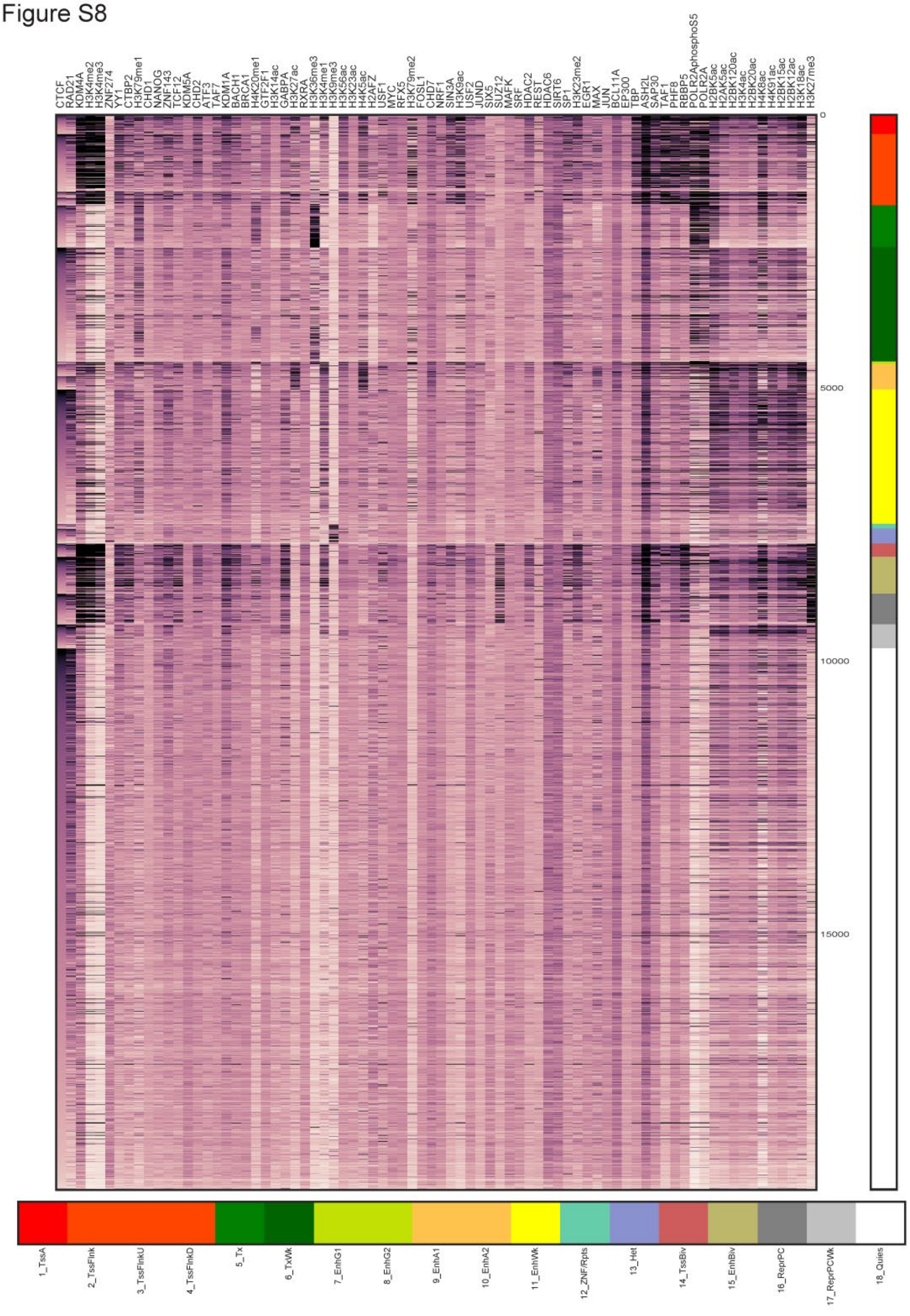

**Fig. S9. CTCF-enriched and -depleted looping interactions**

(A) Distribution of CTCF ChIP-Seq enrichment for all ESC loop anchors, compared to CTCF enrichment for an equal number of loci randomly shifted by distances between 80-160 kb from loop anchors.

(B-C) Chromatin states for CTCF-enriched and -depleted loop anchors, using two different definitions for CTCF enrichment/depletion as indicated.

Figure S9

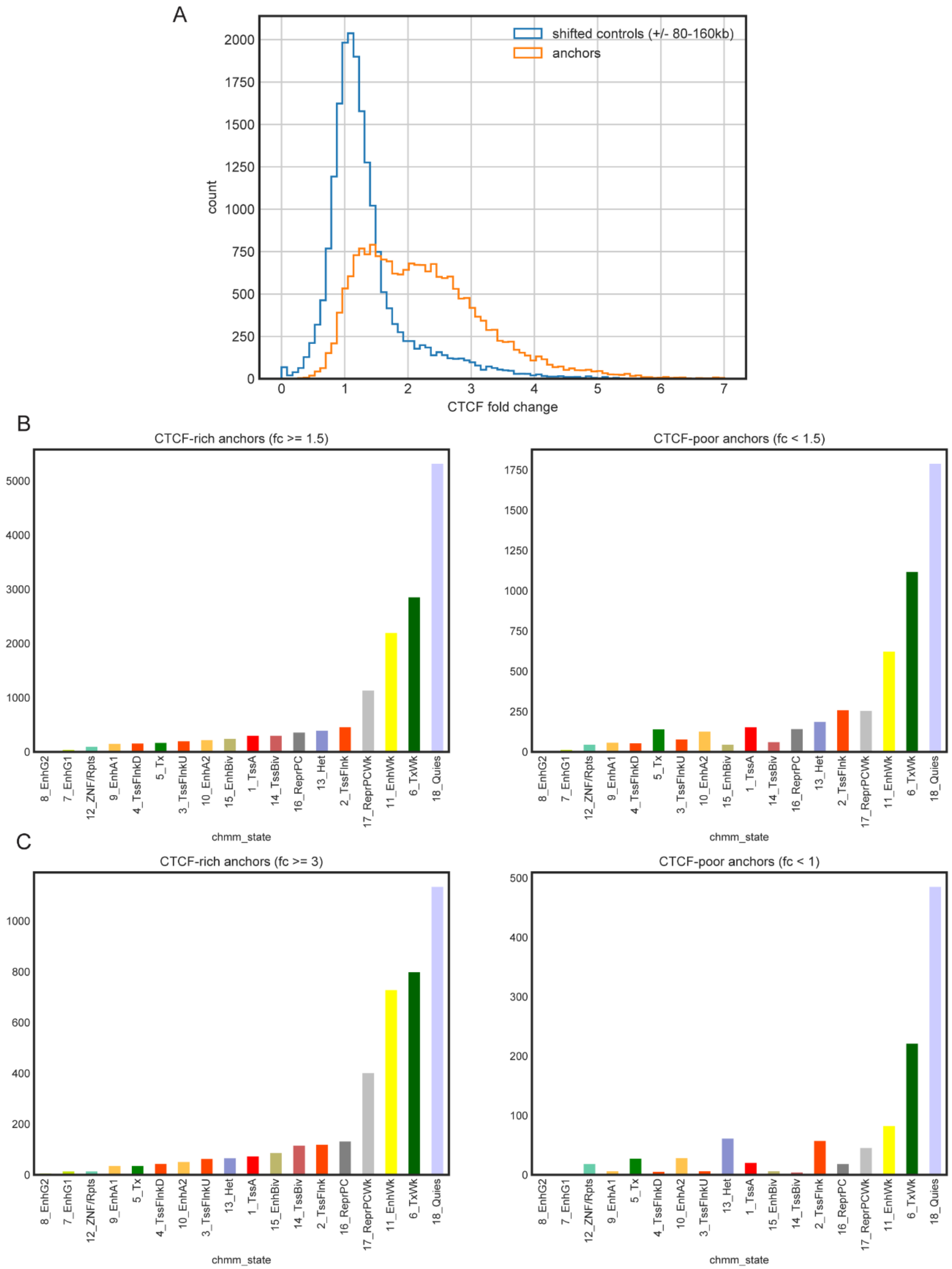

**Fig. S10. Looping interactions at CTCF-depleted loci**

(A) Average contact maps for TSSs (center of x axis) with the nearest enhancer, in both cases excluding regulatory elements overlapping with a significant CTCF ChIP-Seq peak. TSSs are sorted into quintiles based on downstream gene mRNA abundance.

(B) Loops between paired sites for various structural proteins and histone marks. In all cases, we first excluded ChIP peaks for these factors if they fell within 10 kb of a CTCF peak. From this set of peaks, heatmaps show averaged signal for peak pairs falling farther than 5 kb of one another. Red labels indicate factors with <1000 peaks in this analysis.

Figure S10

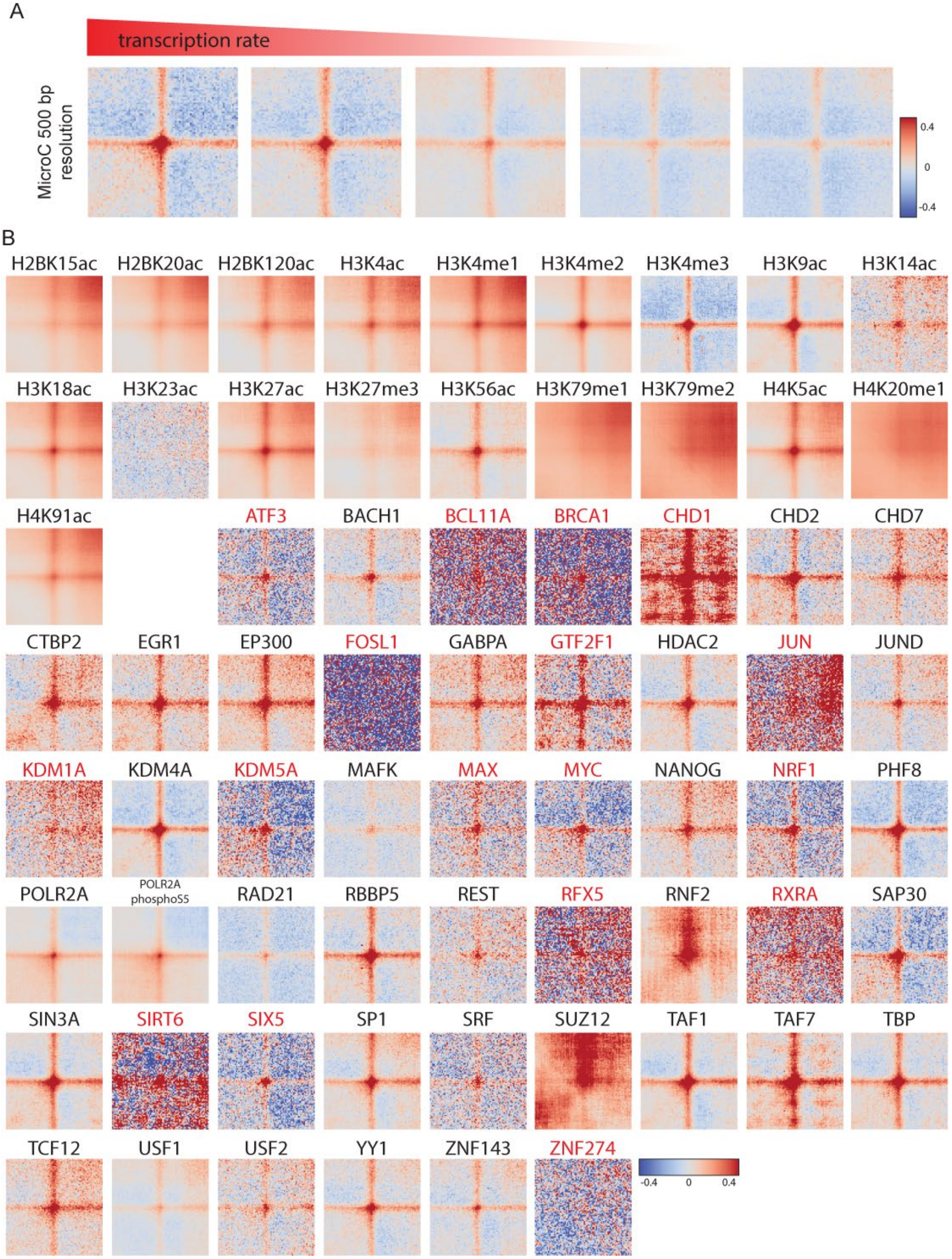
A
transcription rate
MicroC 500 bp resolution
0.4
0
-0.4
B
H2BK15ac
H2BK20ac
H2BK120ac
H3K4ac
H3K4me1
H3K4me2
H3K4me3
H3K9ac
H3K14ac
H3K18ac
H3K23ac
H3K27ac
H3K27me3
H3K56ac
H3K79me1
H3K79me2
H4K5ac
H4K20me1
H4K91ac
ATF3
BACH1
BCL11A
BRCA1
CHD1
CHD2
CHD7
CTBP2
EGR1
EP300
FOSL1
GABPA
GTF2F1
HDAC2
JUN
JUND
KDM1A
KDM4A
KDM5A
MAFK
MAX
MYC
NANOG
NRF1
PHF8
POLR2A
POLR2A phosphoS5
RAD21
RBBP5
REST
RFX5
RNF2
RXRA
SAP30
SIN3A
SIRT6
SIX5
SP1
SRF
SUZ12
TAF1
TAF7
TBP
TCF12
USF1
USF2
YY1
ZNF143
ZNF274
-0.4
0
0.4

**Fig. S11. Loop anchors participate in multiple looping interactions**
Data for loop anchor valency (**Fig. 4F**) are shown here, but with total number of loop anchors (y axis) rather than the fraction of all loop anchors as shown in that figure.

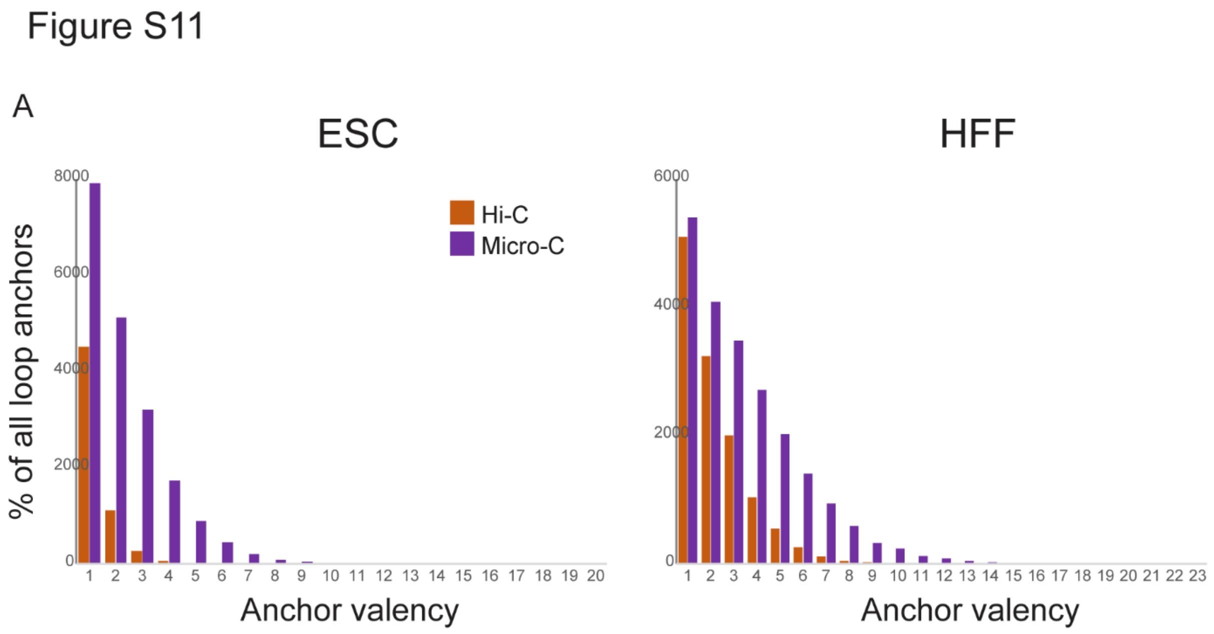

**Table S1.**
Read depth for all four final datasets.

| | HIC - ESC | UC - ESC | HIC - HFF | UC - HFF |
|---|---|---|---|---|
| TOTAL | 6357296696 | 5891219189 | 6477596420 | 7290410596 |
| MAPPED - NO DUP | 2609151200 | 3231252720 | 3018221386 | 4365665012 |
| CIS | 1258316457 | 2793991735 | 2092572184 | 3702356785 |
| TRANS | 1350834743 | 437260985 | 925649202 | 663308227 |

**Table S2.**
Public ChIP-Seq signal datasets used for comparisons to Micro-C 1D heatmaps and composite plots.

| *Experiment.target* | *File.accession* | *Assay* | *Output.type* | *File.format* |
|---|---|---|---|---|
| *RFX5-human* | ENCFF330UHX | ChIP-seq | signal p-value | bigWig |
| *MAFK-human* | ENCFF436UWE | ChIP-seq | signal p-value | bigWig |
| *SRF-human* | ENCFF257UEB | ChIP-seq | signal p-value | bigWig |
| *USF2-human* | ENCFF413HYI | ChIP-seq | signal p-value | bigWig |
| *REST-human* | ENCFF696JDC | ChIP-seq | signal p-value | bigWig |
| *TBP-human* | ENCFF124WHQ | ChIP-seq | signal p-value | bigWig |
| *YY1-human* | ENCFF371JOS | ChIP-seq | signal p-value | bigWig |
| *USF1-human* | ENCFF516AHS | ChIP-seq | signal p-value | bigWig |
| *ZNF143-human* | ENCFF667YWV | ChIP-seq | signal p-value | bigWig |
| *SP1-human* | ENCFF703VUT | ChIP-seq | signal p-value | bigWig |
| *SUZ12-human* | ENCFF609MMJ | ChIP-seq | signal p-value | bigWig |
| *CHD2-human* | ENCFF171BHH | ChIP-seq | signal p-value | bigWig |
| *PHF8-human* | ENCFF378JYM | ChIP-seq | signal p-value | bigWig |
| *EGR1-human* | ENCFF559URO | ChIP-seq | signal p-value | bigWig |
| *JUND-human* | ENCFF189XFG | ChIP-seq | signal p-value | bigWig |
| *SIN3A-human* | ENCFF174DEX | ChIP-seq | signal p-value | bigWig |
| *NRF1-human* | ENCFF952UTI | ChIP-seq | signal p-value | bigWig |

| | | | | |
|---|---|---|---|---|
| *KDM1A-human* | ENCFF297JUX | ChIP-seq | signal p-value | bigWig |
| *CHD1-human* | ENCFF909MSA | ChIP-seq | signal p-value | bigWig |
| *MYC-human* | ENCFF462NJD | ChIP-seq | signal p-value | bigWig |
| *RBBP5-human* | ENCFF696PQE | ChIP-seq | signal p-value | bigWig |
| *KDM5A-human* | ENCFF702KXJ | ChIP-seq | signal p-value | bigWig |
| *BACH1-human* | ENCFF177UEN | ChIP-seq | signal p-value | bigWig |
| *TAF7-human* | ENCFF824JON | ChIP-seq | signal p-value | bigWig |
| *RAD21-human* | ENCFF636BIP | ChIP-seq | signal p-value | bigWig |
| *BRCA1-human* | ENCFF998LWF | ChIP-seq | signal p-value | bigWig |
| *GTF2F1-human* | ENCFF024EFB | ChIP-seq | signal p-value | bigWig |
| *HDAC2-human* | ENCFF723EAE | ChIP-seq | signal p-value | bigWig |
| *SIX5-human* | ENCFF787UNR | ChIP-seq | signal p-value | bigWig |
| *CTCF-human* | ENCFF147GRN | ChIP-seq | signal p-value | bigWig |
| *H3K4me3-human* | ENCFF149EKJ | ChIP-seq | signal p-value | bigWig |
| *POLR2AphosphoS5-human* | ENCFF834EMW | ChIP-seq | signal p-value | bigWig |
| *H3K27ac-human* | ENCFF455PUU | ChIP-seq | signal p-value | bigWig |
| *TAF1-human* | ENCFF569TPM | ChIP-seq | signal p-value | bigWig |
| *H3K14ac-human* | ENCFF394VNN | ChIP-seq | signal p-value | bigWig |
| *H2BK120ac-human* | ENCFF743BTT | ChIP-seq | signal p-value | bigWig |
| *H2BK20ac-human* | ENCFF968MBB | ChIP-seq | signal p-value | bigWig |

| | | | | |
|---|---|---|---|---|
| *H3K23me2-human* | ENCFF541FMI | ChIP-seq | signal p-value | bigWig |
| *H2BK12ac-human* | ENCFF724QZZ | ChIP-seq | signal p-value | bigWig |
| *H2AK5ac-human* | ENCFF195GFI | ChIP-seq | signal p-value | bigWig |
| *H3K9ac-human* | ENCFF392PJM | ChIP-seq | signal p-value | bigWig |
| *H2BK5ac-human* | ENCFF803QKI | ChIP-seq | signal p-value | bigWig |
| *CHD7-human* | ENCFF102VHY | ChIP-seq | signal p-value | bigWig |
| *EP300-human* | ENCFF800HRV | ChIP-seq | signal p-value | bigWig |
| *POLR2A-human* | ENCFF752OKK | ChIP-seq | signal p-value | bigWig |
| *H3K23ac-human* | ENCFF525YNS | ChIP-seq | signal p-value | bigWig |
| *H3K4ac-human* | ENCFF183DXB | ChIP-seq | signal p-value | bigWig |
| *H3K79me1-human* | ENCFF609YPN | ChIP-seq | signal p-value | bigWig |
| *H3K56ac-human* | ENCFF135ZAJ | ChIP-seq | signal p-value | bigWig |
| *H2BK15ac-human* | ENCFF344DLU | ChIP-seq | signal p-value | bigWig |
| *H4K91ac-human* | ENCFF063HYF | ChIP-seq | signal p-value | bigWig |
| *H4K5ac-human* | ENCFF940DMX | ChIP-seq | signal p-value | bigWig |
| *JUN-human* | ENCFF625MXI | ChIP-seq | signal p-value | bigWig |
| *ASH2L-human* | ENCFF210YFI | ChIP-seq | signal p-value | bigWig |

**Table S3.**
Public ChIP-Seq peak position datasets used for Looping interactions pile-ups at CTCF-depleted loci

| *Experiment.target* | *File.accession* | *Assay* | *Output.type* | *File.format* |
|---|---|---|---|---|

| | | | | |
|---|---|---|---|---|
| *ATF3-human* | ENCFF487GLV | ChIP-seq | optimal idr thresholded peaks | bed narrowPeak |
| *BACH1-human* | ENCFF851YHG | ChIP-seq | optimal idr thresholded peaks | bed narrowPeak |
| *BCL11A-human* | ENCFF533KIC | ChIP-seq | pseudoreplicated idr thresholded peaks | bed narrowPeak |
| *BRCA1-human* | ENCFF962YTC | ChIP-seq | optimal idr thresholded peaks | bed narrowPeak |
| *CHD1-human* | ENCFF806HXY | ChIP-seq | optimal idr thresholded peaks | bed narrowPeak |
| *CHD2-human* | ENCFF245UXM | ChIP-seq | optimal idr thresholded peaks | bed narrowPeak |
| *CHD7-human* | ENCFF658SXI | ChIP-seq | optimal idr thresholded peaks | bed narrowPeak |
| *CTBP2-human* | ENCFF482FLE | ChIP-seq | optimal idr thresholded peaks | bed narrowPeak |
| *CTCF-human* | ENCFF821AQO | ChIP-seq | optimal idr thresholded peaks | bed narrowPeak |
| *EGR1-human* | ENCFF477ANT | ChIP-seq | optimal idr thresholded peaks | bed narrowPeak |
| *EP300-human* | ENCFF834UVX | ChIP-seq | optimal idr thresholded peaks | bed narrowPeak |
| *FOSL1-human* | ENCFF063OKB | ChIP-seq | optimal idr thresholded peaks | bed narrowPeak |
| *GABPA-human* | ENCFF225GFQ | ChIP-seq | optimal idr thresholded peaks | bed narrowPeak |
| *GTF2F1-human* | ENCFF493PRB | ChIP-seq | optimal idr thresholded peaks | bed narrowPeak |

| | | | | |
|---|---|---|---|---|
| *H2AFZ-human* | ENCFF745GKP | ChIP-seq | replicated peaks | bed narrowPeak |
| *H2AK5ac-human* | ENCFF160CEI | ChIP-seq | replicated peaks | bed narrowPeak |
| *H2BK120ac-human* | ENCFF808BIC | ChIP-seq | replicated peaks | bed narrowPeak |
| *H2BK12ac-human* | ENCFF315NAV | ChIP-seq | replicated peaks | bed narrowPeak |
| *H2BK15ac-human* | ENCFF997ZDN | ChIP-seq | replicated peaks | bed narrowPeak |
| *H2BK20ac-human* | ENCFF628KOM | ChIP-seq | replicated peaks | bed narrowPeak |
| *H2BK5ac-human* | ENCFF068IZP | ChIP-seq | replicated peaks | bed narrowPeak |
| *H3K14ac-human* | ENCFF642IBN | ChIP-seq | replicated peaks | bed narrowPeak |
| *H3K18ac-human* | ENCFF397EFT | ChIP-seq | replicated peaks | bed narrowPeak |
| *H3K23ac-human* | ENCFF851FNE | ChIP-seq | replicated peaks | bed narrowPeak |
| *H3K27ac-human* | ENCFF045CUG | ChIP-seq | replicated peaks | bed narrowPeak |
| *H3K27me3-human* | ENCFF411ESN | ChIP-seq | replicated peaks | bed narrowPeak |
| *H3K4ac-human* | ENCFF711LQB | ChIP-seq | replicated peaks | bed narrowPeak |
| *H3K4me1-human* | ENCFF558IKG | ChIP-seq | replicated peaks | bed narrowPeak |
| *H3K4me2-human* | ENCFF441MSJ | ChIP-seq | replicated peaks | bed narrowPeak |
| *H3K4me3-human* | ENCFF744ORJ | ChIP-seq | replicated peaks | bed narrowPeak |
| *H3K56ac-human* | ENCFF053HKF | ChIP-seq | replicated peaks | bed narrowPeak |
| *H3K79me1-human* | ENCFF385DSV | ChIP-seq | replicated peaks | bed narrowPeak |
| *H3K79me2-human* | ENCFF202EZL | ChIP-seq | replicated peaks | bed narrowPeak |
| *H3K9ac-human* | ENCFF343GTP | ChIP-seq | replicated peaks | bed narrowPeak |

| | | | | |
|---|---|---|---|---|
| *H3K9me3-human* | ENCFF250GSY | ChIP-seq | replicated peaks | bed narrowPeak |
| *H4K20me1-human* | ENCFF439CWL | ChIP-seq | replicated peaks | bed narrowPeak |
| *H4K20me1-human* | ENCFF997CKL | ChIP-seq | replicated peaks | bed narrowPeak |
| *H4K5ac-human* | ENCFF237OAY | ChIP-seq | replicated peaks | bed narrowPeak |
| *H4K8ac-human* | ENCFF760EFQ | ChIP-seq | replicated peaks | bed narrowPeak |
| *H4K91ac-human* | ENCFF964FVB | ChIP-seq | replicated peaks | bed narrowPeak |
| *HDAC2-human* | ENCFF497YNJ | ChIP-seq | optimal idr thresholded peaks | bed narrowPeak |
| *JUN-human* | ENCFF312GEN | ChIP-seq | optimal idr thresholded peaks | bed narrowPeak |
| *JUND-human* | ENCFF443HNU | ChIP-seq | optimal idr thresholded peaks | bed narrowPeak |
| *KDM1A-human* | ENCFF562OAN | ChIP-seq | optimal idr thresholded peaks | bed narrowPeak |
| *KDM4A-human* | ENCFF205WRX | ChIP-seq | pseudoreplicated idr thresholded peaks | bed narrowPeak |
| *KDM5A-human* | ENCFF342EEV | ChIP-seq | optimal idr thresholded peaks | bed narrowPeak |
| *MAFK-human* | ENCFF712RIS | ChIP-seq | optimal idr thresholded peaks | bed narrowPeak |
| *MAX-human* | ENCFF762LKG | ChIP-seq | optimal idr thresholded peaks | bed narrowPeak |
| *MYC-human* | ENCFF392JJN | ChIP-seq | optimal idr thresholded peaks | bed narrowPeak |
| *NANOG-human* | ENCFF794GVQ | ChIP-seq | optimal idr thresholded peaks | bed narrowPeak |

| | | | | |
|---|---|---|---|---|
| *NRF1-human* | ENCFF407IVS | ChIP-seq | optimal idr thresholded peaks | bed narrowPeak |
| *PHF8-human* | ENCFF651QOL | ChIP-seq | optimal idr thresholded peaks | bed narrowPeak |
| *POLR2A-human* | ENCFF422HDN | ChIP-seq | optimal idr thresholded peaks | bed narrowPeak |
| *POLR2AphosphoS5-human* | ENCFF418QVJ | ChIP-seq | optimal idr thresholded peaks | bed narrowPeak |
| *RAD21-human* | ENCFF255FRL | ChIP-seq | optimal idr thresholded peaks | bed narrowPeak |
| *RBBP5-human* | ENCFF607WCG | ChIP-seq | optimal idr thresholded peaks | bed narrowPeak |
| *REST-human* | ENCFF779CWH | ChIP-seq | optimal idr thresholded peaks | bed narrowPeak |
| *RFX5-human* | ENCFF062WBN | ChIP-seq | optimal idr thresholded peaks | bed narrowPeak |
| *RNF2-human* | ENCFF283MNG | ChIP-seq | optimal idr thresholded peaks | bed narrowPeak |
| *RXRA-human* | ENCFF430SIE | ChIP-seq | optimal idr thresholded peaks | bed narrowPeak |
| *SAP30-human* | ENCFF193TFR | ChIP-seq | optimal idr thresholded peaks | bed narrowPeak |
| *SIN3A-human* | ENCFF514BGQ | ChIP-seq | optimal idr thresholded peaks | bed narrowPeak |
| *SIRT6-human* | ENCFF539KSF | ChIP-seq | pseudoreplicated idr thresholded peaks | bed narrowPeak |
| *SIX5-human* | ENCFF644BNN | ChIP-seq | optimal idr thresholded peaks | bed narrowPeak |

| | | | | |
|---|---|---|---|---|
| *SP1-human* | ENCFF500JFI | ChIP-seq | optimal idr thresholded peaks | bed narrowPeak |
| *SRF-human* | ENCFF345IDL | ChIP-seq | optimal idr thresholded peaks | bed narrowPeak |
| *SUZ12-human* | ENCFF938PCI | ChIP-seq | optimal idr thresholded peaks | bed narrowPeak |
| *SUZ12-human* | ENCFF671SZQ | ChIP-seq | optimal idr thresholded peaks | bed narrowPeak |
| *TAF1-human* | ENCFF870SFJ | ChIP-seq | optimal idr thresholded peaks | bed narrowPeak |
| *TAF7-human* | ENCFF243PSJ | ChIP-seq | optimal idr thresholded peaks | bed narrowPeak |
| *TBP-human* | ENCFF748YXF | ChIP-seq | optimal idr thresholded peaks | bed narrowPeak |
| *TCF12-human* | ENCFF740HPV | ChIP-seq | optimal idr thresholded peaks | bed narrowPeak |
| *USF1-human* | ENCFF699HXL | ChIP-seq | optimal idr thresholded peaks | bed narrowPeak |
| *USF2-human* | ENCFF710JBU | ChIP-seq | optimal idr thresholded peaks | bed narrowPeak |
| *YY1-human* | ENCFF509GYP | ChIP-seq | optimal idr thresholded peaks | bed narrowPeak |
| *ZNF143-human* | ENCFF933WSP | ChIP-seq | optimal idr thresholded peaks | bed narrowPeak |
| *ZNF274-human* | ENCFF943NOT | ChIP-seq | optimal idr thresholded peaks | bed narrowPeak |

**Table S4.**
Public ChIP-Seq datasets used for Epigenetic characterization of ESC dot anchors.

| *Experiment target* | *File accession* | *Assay* | *Output type* | *File format* |
|---|---|---|---|---|
| *ZNF274-human* | ENCFF160MNX | ChIP-seq | fold change over control | bigWig |
| *ZNF274-human* | ENCFF040IXF | ChIP-seq | fold change over control | bigWig |
| *ZNF274-human* | ENCFF060FUO | ChIP-seq | fold change over control | bigWig |
| *H3K79me2-human* | ENCFF460SAM | ChIP-seq | fold change over control | bigWig |
| *H3K79me2-human* | ENCFF879LBS | ChIP-seq | fold change over control | bigWig |
| *H3K79me2-human* | ENCFF854LTH | ChIP-seq | fold change over control | bigWig |
| *TBP-human* | ENCFF515VTD | ChIP-seq | fold change over control | bigWig |
| *TBP-human* | ENCFF747EIS | ChIP-seq | fold change over control | bigWig |
| *TBP-human* | ENCFF052TRV | ChIP-seq | fold change over control | bigWig |
| *ASH2L-human* | ENCFF806OYX | ChIP-seq | fold change over control | bigWig |
| *FOSL1-human* | ENCFF616GYO | ChIP-seq | fold change over control | bigWig |
| *FOSL1-human* | ENCFF498IQF | ChIP-seq | fold change over control | bigWig |
| *FOSL1-human* | ENCFF413VTX | ChIP-seq | fold change over control | bigWig |
| *CHD7-human* | ENCFF450JZY | ChIP-seq | fold change over control | bigWig |
| *CHD7-human* | ENCFF575OWE | ChIP-seq | fold change over control | bigWig |
| *CHD7-human* | ENCFF530LBH | ChIP-seq | fold change over control | bigWig |
| *NRF1-human* | ENCFF515ERL | ChIP-seq | fold change over control | bigWig |
| *NRF1-human* | ENCFF449KGF | ChIP-seq | fold change over control | bigWig |
| *NRF1-human* | ENCFF152BEY | ChIP-seq | fold change over control | bigWig |
| *SIN3A-human* | ENCFF016TFI | ChIP-seq | fold change over control | bigWig |
| *SIN3A-human* | ENCFF350OAA | ChIP-seq | fold change over control | bigWig |

| | | | | |
|---|---|---|---|---|
| *SIN3A-human* | ENCFF875ANJ | ChIP-seq | fold change over control | bigWig |
| *RAD21-human* | ENCFF904MGK | ChIP-seq | fold change over control | bigWig |
| *RAD21-human* | ENCFF164PQM | ChIP-seq | fold change over control | bigWig |
| *RAD21-human* | ENCFF791XXJ | ChIP-seq | fold change over control | bigWig |
| *H3K9ac-human* | ENCFF758JKT | ChIP-seq | fold change over control | bigWig |
| *H3K9ac-human* | ENCFF807XBN | ChIP-seq | fold change over control | bigWig |
| *H3K9ac-human* | ENCFF795DQH | ChIP-seq | fold change over control | bigWig |
| *USF2-human* | ENCFF372HIM | ChIP-seq | fold change over control | bigWig |
| *USF2-human* | ENCFF757FPX | ChIP-seq | fold change over control | bigWig |
| *USF2-human* | ENCFF870KXV | ChIP-seq | fold change over control | bigWig |
| *HDAC2-human* | ENCFF192XRQ | ChIP-seq | fold change over control | bigWig |
| *HDAC2-human* | ENCFF290ALW | ChIP-seq | fold change over control | bigWig |
| *HDAC2-human* | ENCFF948IYF | ChIP-seq | fold change over control | bigWig |
| *SIX5-human* | ENCFF737SHB | ChIP-seq | fold change over control | bigWig |
| *SIX5-human* | ENCFF665USC | ChIP-seq | fold change over control | bigWig |
| *SIX5-human* | ENCFF028IYH | ChIP-seq | fold change over control | bigWig |
| *PHF8-human* | ENCFF401PWV | ChIP-seq | fold change over control | bigWig |
| *PHF8-human* | ENCFF888QPY | ChIP-seq | fold change over control | bigWig |
| *PHF8-human* | ENCFF935JRI | ChIP-seq | fold change over control | bigWig |
| *SUZ12-human* | ENCFF693IMS | ChIP-seq | fold change over control | bigWig |
| *SUZ12-human* | ENCFF429YPI | ChIP-seq | fold change over control | bigWig |
| *SUZ12-human* | ENCFF008SLD | ChIP-seq | fold change over control | bigWig |
| *MAFK-human* | ENCFF637DVH | ChIP-seq | fold change over control | bigWig |
| *MAFK-human* | ENCFF640RNH | ChIP-seq | fold change over control | bigWig |

| | | | | |
|---|---|---|---|---|
| *MAFK-human* | ENCFF407RRX | ChIP-seq | fold change over control | bigWig |
| *HDAC2-human* | ENCFF424DKN | ChIP-seq | fold change over control | bigWig |
| *HDAC2-human* | ENCFF593FGQ | ChIP-seq | fold change over control | bigWig |
| *HDAC2-human* | ENCFF640QBB | ChIP-seq | fold change over control | bigWig |
| *SRF-human* | ENCFF996WRR | ChIP-seq | fold change over control | bigWig |
| *SRF-human* | ENCFF556LAR | ChIP-seq | fold change over control | bigWig |
| *SRF-human* | ENCFF941KEV | ChIP-seq | fold change over control | bigWig |
| *EP300-human* | ENCFF489CQP | ChIP-seq | fold change over control | bigWig |
| *HDAC6-human* | ENCFF243FNC | ChIP-seq | fold change over control | bigWig |
| *H3K4me3-human* | ENCFF042KLQ | ChIP-seq | fold change over control | bigWig |
| *H3K4me3-human* | ENCFF525RSM | ChIP-seq | fold change over control | bigWig |
| *H3K4me3-human* | ENCFF629XVB | ChIP-seq | fold change over control | bigWig |
| *H2BK20ac-human* | ENCFF592YSB | ChIP-seq | fold change over control | bigWig |
| *H2BK20ac-human* | ENCFF250ZWC | ChIP-seq | fold change over control | bigWig |
| *H2BK20ac-human* | ENCFF382GFP | ChIP-seq | fold change over control | bigWig |
| *H3K27me3-human* | ENCFF819LHK | ChIP-seq | fold change over control | bigWig |
| *H3K27me3-human* | ENCFF371XFA | ChIP-seq | fold change over control | bigWig |
| *H3K27me3-human* | ENCFF095OPQ | ChIP-seq | fold change over control | bigWig |
| *H3K23me2-human* | ENCFF517UOA | ChIP-seq | fold change over control | bigWig |
| *H3K23me2-human* | ENCFF074ALX | ChIP-seq | fold change over control | bigWig |
| *H3K23me2-human* | ENCFF757RMF | ChIP-seq | fold change over control | bigWig |
| *H3K27me3-human* | ENCFF842IJO | ChIP-seq | fold change over control | bigWig |
| *H3K27me3-human* | ENCFF912ZUR | ChIP-seq | fold change over control | bigWig |
| *H3K27me3-human* | ENCFF976VVR | ChIP-seq | fold change over control | bigWig |

| | | | | |
|---|---|---|---|---|
| *EGR1-human* | ENCFF621LNV | ChIP-seq | fold change over control | bigWig |
| *EGR1-human* | ENCFF464FML | ChIP-seq | fold change over control | bigWig |
| *EGR1-human* | ENCFF341OGJ | ChIP-seq | fold change over control | bigWig |
| *MAX-human* | ENCFF757VYL | ChIP-seq | fold change over control | bigWig |
| *MAX-human* | ENCFF857DHC | ChIP-seq | fold change over control | bigWig |
| *MAX-human* | ENCFF444FFZ | ChIP-seq | fold change over control | bigWig |
| *H3K4me3-human* | ENCFF368PQO | ChIP-seq | fold change over control | bigWig |
| *H3K4me3-human* | ENCFF473GLI | ChIP-seq | fold change over control | bigWig |
| *H3K4me3-human* | ENCFF730VVX | ChIP-seq | fold change over control | bigWig |
| *SUZ12-human* | ENCFF652TAQ | ChIP-seq | fold change over control | bigWig |
| *SUZ12-human* | ENCFF723MAM | ChIP-seq | fold change over control | bigWig |
| *SUZ12-human* | ENCFF887KVL | ChIP-seq | fold change over control | bigWig |
| *JUN-human* | ENCFF417OBC | ChIP-seq | fold change over control | bigWig |
| *JUN-human* | ENCFF459HBP | ChIP-seq | fold change over control | bigWig |
| *JUN-human* | ENCFF815WEI | ChIP-seq | fold change over control | bigWig |
| *BCL11A-human* | ENCFF203IVF | ChIP-seq | fold change over control | bigWig |
| *RNF2-human* | ENCFF439ACI | ChIP-seq | fold change over control | bigWig |
| *RNF2-human* | ENCFF539IQR | ChIP-seq | fold change over control | bigWig |
| *RNF2-human* | ENCFF308TCO | ChIP-seq | fold change over control | bigWig |
| *KDM4A-human* | ENCFF916KLB | ChIP-seq | fold change over control | bigWig |
| *TAF1-human* | ENCFF894CVE | ChIP-seq | fold change over control | bigWig |
| *TAF1-human* | ENCFF689QWC | ChIP-seq | fold change over control | bigWig |
| *TAF1-human* | ENCFF360PPQ | ChIP-seq | fold change over control | bigWig |
| *REST-human* | ENCFF390YMS | ChIP-seq | fold change over control | bigWig |

| | | | | |
|---|---|---|---|---|
| *REST-human* | ENCFF707FDL | ChIP-seq | fold change over control | bigWig |
| *REST-human* | ENCFF600PQH | ChIP-seq | fold change over control | bigWig |
| *SP1-human* | ENCFF286DIH | ChIP-seq | fold change over control | bigWig |
| *SP1-human* | ENCFF861UXW | ChIP-seq | fold change over control | bigWig |
| *SP1-human* | ENCFF256MVQ | ChIP-seq | fold change over control | bigWig |
| *SIRT6-human* | ENCFF822ZVD | ChIP-seq | fold change over control | bigWig |
| *RFX5-human* | ENCFF502KGA | ChIP-seq | fold change over control | bigWig |
| *RFX5-human* | ENCFF279KGJ | ChIP-seq | fold change over control | bigWig |
| *RFX5-human* | ENCFF027CMH | ChIP-seq | fold change over control | bigWig |
| *JUND-human* | ENCFF220AJW | ChIP-seq | fold change over control | bigWig |
| *JUND-human* | ENCFF339LKC | ChIP-seq | fold change over control | bigWig |
| *JUND-human* | ENCFF744ISC | ChIP-seq | fold change over control | bigWig |
| *USF1-human* | ENCFF511BGE | ChIP-seq | fold change over control | bigWig |
| *USF1-human* | ENCFF708QRS | ChIP-seq | fold change over control | bigWig |
| *USF1-human* | ENCFF133IZI | ChIP-seq | fold change over control | bigWig |
| *KDM1A-human* | ENCFF272TTV | ChIP-seq | fold change over control | bigWig |
| *KDM1A-human* | ENCFF563SQA | ChIP-seq | fold change over control | bigWig |
| *KDM1A-human* | ENCFF222RPJ | ChIP-seq | fold change over control | bigWig |
| *YY1-human* | ENCFF515UYM | ChIP-seq | fold change over control | bigWig |
| *YY1-human* | ENCFF170XRZ | ChIP-seq | fold change over control | bigWig |
| *YY1-human* | ENCFF406PYH | ChIP-seq | fold change over control | bigWig |
| *CTCF-human* | ENCFF586WDJ | ChIP-seq | fold change over control | bigWig |
| *CTBP2-human* | ENCFF231ZHA | ChIP-seq | fold change over control | bigWig |
| *CTBP2-human* | ENCFF479EQH | ChIP-seq | fold change over control | bigWig |

| | | | | |
|---|---|---|---|---|
| *CTBP2-human* | ENCFF562PRB | ChIP-seq | fold change over control | bigWig |
| *SIN3A-human* | ENCFF503MAL | ChIP-seq | fold change over control | bigWig |
| *SIN3A-human* | ENCFF571JNW | ChIP-seq | fold change over control | bigWig |
| *SIN3A-human* | ENCFF080NJX | ChIP-seq | fold change over control | bigWig |
| *MYC-human* | ENCFF865USR | ChIP-seq | fold change over control | bigWig |
| *MYC-human* | ENCFF491NAO | ChIP-seq | fold change over control | bigWig |
| *MYC-human* | ENCFF878ZLF | ChIP-seq | fold change over control | bigWig |
| *JUND-human* | ENCFF655OWT | ChIP-seq | fold change over control | bigWig |
| *JUND-human* | ENCFF790OJE | ChIP-seq | fold change over control | bigWig |
| *JUND-human* | ENCFF128BVN | ChIP-seq | fold change over control | bigWig |
| *CHD1-human* | ENCFF452YHL | ChIP-seq | fold change over control | bigWig |
| *CHD1-human* | ENCFF926LHR | ChIP-seq | fold change over control | bigWig |
| *CHD1-human* | ENCFF576FUZ | ChIP-seq | fold change over control | bigWig |
| *NANOG-human* | ENCFF479ERO | ChIP-seq | fold change over control | bigWig |
| *NANOG-human* | ENCFF305LHR | ChIP-seq | fold change over control | bigWig |
| *NANOG-human* | ENCFF082LGX | ChIP-seq | fold change over control | bigWig |
| *H3K4me2-human* | ENCFF571WYL | ChIP-seq | fold change over control | bigWig |
| *H3K4me2-human* | ENCFF142UOO | ChIP-seq | fold change over control | bigWig |
| *H3K4me2-human* | ENCFF903RZE | ChIP-seq | fold change over control | bigWig |
| *ZNF143-human* | ENCFF025QWP | ChIP-seq | fold change over control | bigWig |
| *ZNF143-human* | ENCFF480ASE | ChIP-seq | fold change over control | bigWig |
| *ZNF143-human* | ENCFF377SDG | ChIP-seq | fold change over control | bigWig |
| *TCF12-human* | ENCFF143RCD | ChIP-seq | fold change over control | bigWig |
| *TCF12-human* | ENCFF548GTM | ChIP-seq | fold change over control | bigWig |

| | | | | |
|---|---|---|---|---|
| *TCF12-human* | ENCFF715MYQ | ChIP-seq | fold change over control | bigWig |
| *KDM5A-human* | ENCFF708KOI | ChIP-seq | fold change over control | bigWig |
| *KDM5A-human* | ENCFF825WLX | ChIP-seq | fold change over control | bigWig |
| *KDM5A-human* | ENCFF006CJS | ChIP-seq | fold change over control | bigWig |
| *CHD2-human* | ENCFF977BAU | ChIP-seq | fold change over control | bigWig |
| *CHD2-human* | ENCFF509LLY | ChIP-seq | fold change over control | bigWig |
| *CHD2-human* | ENCFF318NSO | ChIP-seq | fold change over control | bigWig |
| *RBBP5-human* | ENCFF485HWP | ChIP-seq | fold change over control | bigWig |
| *RBBP5-human* | ENCFF692MWE | ChIP-seq | fold change over control | bigWig |
| *RBBP5-human* | ENCFF076ZMU | ChIP-seq | fold change over control | bigWig |
| *ATF3-human* | ENCFF767IYZ | ChIP-seq | fold change over control | bigWig |
| *ATF3-human* | ENCFF481EHX | ChIP-seq | fold change over control | bigWig |
| *ATF3-human* | ENCFF889EVL | ChIP-seq | fold change over control | bigWig |
| *TAF7-human* | ENCFF777AYI | ChIP-seq | fold change over control | bigWig |
| *TAF7-human* | ENCFF195SPX | ChIP-seq | fold change over control | bigWig |
| *TAF7-human* | ENCFF160JKQ | ChIP-seq | fold change over control | bigWig |
| *POLR2A-human* | ENCFF975VIV | ChIP-seq | fold change over control | bigWig |
| *SAP30-human* | ENCFF240MHG | ChIP-seq | fold change over control | bigWig |
| *SAP30-human* | ENCFF459CAJ | ChIP-seq | fold change over control | bigWig |
| *SAP30-human* | ENCFF779YFX | ChIP-seq | fold change over control | bigWig |
| *BACH1-human* | ENCFF872ZCR | ChIP-seq | fold change over control | bigWig |
| *BACH1-human* | ENCFF445KLN | ChIP-seq | fold change over control | bigWig |
| *BACH1-human* | ENCFF594ALF | ChIP-seq | fold change over control | bigWig |
| *H2AFZ-human* | ENCFF775ZWT | ChIP-seq | fold change over control | bigWig |

| | | | | |
|---|---|---|---|---|
| *H2AFZ-human* | ENCFF801HJQ | ChIP-seq | fold change over control | bigWig |
| *H2AFZ-human* | ENCFF575EAK | ChIP-seq | fold change over control | bigWig |
| *BCL11A-human* | ENCFF735TIA | ChIP-seq | fold change over control | bigWig |
| *CTCF-human* | ENCFF520THR | ChIP-seq | fold change over control | bigWig |
| *REST-human* | ENCFF533SBV | ChIP-seq | fold change over control | bigWig |
| *REST-human* | ENCFF648WCI | ChIP-seq | fold change over control | bigWig |
| *REST-human* | ENCFF833NSS | ChIP-seq | fold change over control | bigWig |
| *RAD21-human* | ENCFF182WAH | ChIP-seq | fold change over control | bigWig |
| *RAD21-human* | ENCFF130VDU | ChIP-seq | fold change over control | bigWig |
| *RAD21-human* | ENCFF913JGA | ChIP-seq | fold change over control | bigWig |
| *CTCF-human* | ENCFF549TOE | ChIP-seq | fold change over control | bigWig |
| *CTCF-human* | ENCFF493IMI | ChIP-seq | fold change over control | bigWig |
| *CTCF-human* | ENCFF473IZV | ChIP-seq | fold change over control | bigWig |
| *BRCA1-human* | ENCFF971FXL | ChIP-seq | fold change over control | bigWig |
| *BRCA1-human* | ENCFF620MRE | ChIP-seq | fold change over control | bigWig |
| *BRCA1-human* | ENCFF002NDR | ChIP-seq | fold change over control | bigWig |
| *H4K20me1-human* | ENCFF026MPU | ChIP-seq | fold change over control | bigWig |
| *H4K20me1-human* | ENCFF861LCZ | ChIP-seq | fold change over control | bigWig |
| *H4K20me1-human* | ENCFF067XIV | ChIP-seq | fold change over control | bigWig |
| *GTF2F1-human* | ENCFF454ZYL | ChIP-seq | fold change over control | bigWig |
| *GTF2F1-human* | ENCFF173BEC | ChIP-seq | fold change over control | bigWig |
| *GTF2F1-human* | ENCFF500KUC | ChIP-seq | fold change over control | bigWig |
| *H3K14ac-human* | ENCFF983XBZ | ChIP-seq | fold change over control | bigWig |
| *H3K14ac-human* | ENCFF809NBO | ChIP-seq | fold change over control | bigWig |

| | | | | |
|---|---|---|---|---|
| *H3K14ac-human* | ENCFF605ROH | ChIP-seq | fold change over control | bigWig |
| *GABPA-human* | ENCFF542BRR | ChIP-seq | fold change over control | bigWig |
| *GABPA-human* | ENCFF931OAX | ChIP-seq | fold change over control | bigWig |
| *GABPA-human* | ENCFF401DOJ | ChIP-seq | fold change over control | bigWig |
| *H3K79me2-human* | ENCFF798HCY | ChIP-seq | fold change over control | bigWig |
| *H3K79me2-human* | ENCFF833AVU | ChIP-seq | fold change over control | bigWig |
| *H3K79me2-human* | ENCFF234WMO | ChIP-seq | fold change over control | bigWig |
| *H3K27ac-human* | ENCFF872FNR | ChIP-seq | fold change over control | bigWig |
| *H3K27ac-human* | ENCFF671QJO | ChIP-seq | fold change over control | bigWig |
| *H3K27ac-human* | ENCFF986PCY | ChIP-seq | fold change over control | bigWig |
| *RXRA-human* | ENCFF134SMY | ChIP-seq | fold change over control | bigWig |
| *RXRA-human* | ENCFF793PJS | ChIP-seq | fold change over control | bigWig |
| *RXRA-human* | ENCFF634SQG | ChIP-seq | fold change over control | bigWig |
| *H3K36me3-human* | ENCFF510INK | ChIP-seq | fold change over control | bigWig |
| *H3K36me3-human* | ENCFF637IPJ | ChIP-seq | fold change over control | bigWig |
| *H3K36me3-human* | ENCFF422PZQ | ChIP-seq | fold change over control | bigWig |
| *CHD1-human* | ENCFF708JWR | ChIP-seq | fold change over control | bigWig |
| *CHD1-human* | ENCFF193UAD | ChIP-seq | fold change over control | bigWig |
| *CHD1-human* | ENCFF563QHP | ChIP-seq | fold change over control | bigWig |
| *H2BK12ac-human* | ENCFF342ILQ | ChIP-seq | fold change over control | bigWig |
| *H2BK12ac-human* | ENCFF949KSH | ChIP-seq | fold change over control | bigWig |
| *H2BK12ac-human* | ENCFF873OYG | ChIP-seq | fold change over control | bigWig |
| *H3K9ac-human* | ENCFF605BDO | ChIP-seq | fold change over control | bigWig |
| *H3K27me3-human* | ENCFF222VOF | ChIP-seq | fold change over control | bigWig |

| | | | | |
|---|---|---|---|---|
| *H3K27me3-human* | ENCFF043JFV | ChIP-seq | fold change over control | bigWig |
| *H3K27me3-human* | ENCFF822QTE | ChIP-seq | fold change over control | bigWig |
| *H3K36me3-human* | ENCFF138NDD | ChIP-seq | fold change over control | bigWig |
| *H3K36me3-human* | ENCFF261QJO | ChIP-seq | fold change over control | bigWig |
| *H3K36me3-human* | ENCFF918TJS | ChIP-seq | fold change over control | bigWig |
| *H3K36me3-human* | ENCFF209IAH | ChIP-seq | fold change over control | bigWig |
| *H3K36me3-human* | ENCFF526KDV | ChIP-seq | fold change over control | bigWig |
| *H3K36me3-human* | ENCFF591LCS | ChIP-seq | fold change over control | bigWig |
| *H4K20me1-human* | ENCFF772CZB | ChIP-seq | fold change over control | bigWig |
| *H4K20me1-human* | ENCFF333TAK | ChIP-seq | fold change over control | bigWig |
| *H4K20me1-human* | ENCFF511ZNC | ChIP-seq | fold change over control | bigWig |
| *H3K36me3-human* | ENCFF505NMF | ChIP-seq | fold change over control | bigWig |
| *H3K36me3-human* | ENCFF141YAA | ChIP-seq | fold change over control | bigWig |
| *H3K36me3-human* | ENCFF124FMF | ChIP-seq | fold change over control | bigWig |
| *H3K9ac-human* | ENCFF993KIB | ChIP-seq | fold change over control | bigWig |
| *H3K9ac-human* | ENCFF357DHN | ChIP-seq | fold change over control | bigWig |
| *H3K9ac-human* | ENCFF834AZA | ChIP-seq | fold change over control | bigWig |
| *H2BK5ac-human* | ENCFF451CYN | ChIP-seq | fold change over control | bigWig |
| *H2BK5ac-human* | ENCFF692LJR | ChIP-seq | fold change over control | bigWig |
| *H2BK5ac-human* | ENCFF411OPM | ChIP-seq | fold change over control | bigWig |
| *H3K4me1-human* | ENCFF770GBO | ChIP-seq | fold change over control | bigWig |
| *H3K4me1-human* | ENCFF922GHX | ChIP-seq | fold change over control | bigWig |
| *H3K4me1-human* | ENCFF584AVI | ChIP-seq | fold change over control | bigWig |
| *H3K4me2-human* | ENCFF900PRJ | ChIP-seq | fold change over control | bigWig |

| | | | | |
|---|---|---|---|---|
| *H3K4me2-human* | ENCFF502TJG | ChIP-seq | fold change over control | bigWig |
| *H3K4me2-human* | ENCFF713WKN | ChIP-seq | fold change over control | bigWig |
| *H2AK5ac-human* | ENCFF783IVG | ChIP-seq | fold change over control | bigWig |
| *H2AK5ac-human* | ENCFF508WLD | ChIP-seq | fold change over control | bigWig |
| *H2AK5ac-human* | ENCFF909XHX | ChIP-seq | fold change over control | bigWig |
| *POLR2AphosphoS5-human* | ENCFF342XVZ | ChIP-seq | fold change over control | bigWig |
| *POLR2AphosphoS5-human* | ENCFF627MAZ | ChIP-seq | fold change over control | bigWig |
| *POLR2AphosphoS5-human* | ENCFF655OPV | ChIP-seq | fold change over control | bigWig |
| *H3K4me1-human* | ENCFF593OAZ | ChIP-seq | fold change over control | bigWig |
| *H3K4me1-human* | ENCFF618WQF | ChIP-seq | fold change over control | bigWig |
| *H3K4me1-human* | ENCFF031POC | ChIP-seq | fold change over control | bigWig |
| *H3K4me3-human* | ENCFF818FCM | ChIP-seq | fold change over control | bigWig |
| *H3K4me3-human* | ENCFF123HPX | ChIP-seq | fold change over control | bigWig |
| *H3K4me3-human* | ENCFF623ZAW | ChIP-seq | fold change over control | bigWig |
| *H3K9me3-human* | ENCFF206CEF | ChIP-seq | fold change over control | bigWig |
| *H3K9me3-human* | ENCFF917WNU | ChIP-seq | fold change over control | bigWig |
| *H3K9me3-human* | ENCFF752UGN | ChIP-seq | fold change over control | bigWig |
| *POLR2A-human* | ENCFF376XTC | ChIP-seq | fold change over control | bigWig |
| *POLR2A-human* | ENCFF795YDZ | ChIP-seq | fold change over control | bigWig |
| *POLR2A-human* | ENCFF379IRQ | ChIP-seq | fold change over control | bigWig |
| *H3K9me3-human* | ENCFF407AQX | ChIP-seq | fold change over control | bigWig |
| *H3K9me3-human* | ENCFF435YZW | ChIP-seq | fold change over control | bigWig |
| *H3K9me3-human* | ENCFF492JAC | ChIP-seq | fold change over control | bigWig |
| *H3K9me3-human* | ENCFF967ZNE | ChIP-seq | fold change over control | bigWig |

| | | | | |
|---|---|---|---|---|
| *H3K9me3-human* | ENCFF300CCI | ChIP-seq | fold change over control | bigWig |
| *H3K27ac-human* | ENCFF423TVA | ChIP-seq | fold change over control | bigWig |
| *H3K27ac-human* | ENCFF721ZIS | ChIP-seq | fold change over control | bigWig |
| *H3K27ac-human* | ENCFF621NFH | ChIP-seq | fold change over control | bigWig |
| *H3K56ac-human* | ENCFF688YVV | ChIP-seq | fold change over control | bigWig |
| *H3K56ac-human* | ENCFF649JHH | ChIP-seq | fold change over control | bigWig |
| *H3K56ac-human* | ENCFF509ZNJ | ChIP-seq | fold change over control | bigWig |
| *H2BK120ac-human* | ENCFF194TPO | ChIP-seq | fold change over control | bigWig |
| *H2BK120ac-human* | ENCFF757EYT | ChIP-seq | fold change over control | bigWig |
| *H2BK120ac-human* | ENCFF211TVQ | ChIP-seq | fold change over control | bigWig |
| *H3K4ac-human* | ENCFF571UTM | ChIP-seq | fold change over control | bigWig |
| *H3K4ac-human* | ENCFF072LWV | ChIP-seq | fold change over control | bigWig |
| *H3K4ac-human* | ENCFF176INM | ChIP-seq | fold change over control | bigWig |
| *H2AFZ-human* | ENCFF157VMK | ChIP-seq | fold change over control | bigWig |
| *H3K27me3-human* | ENCFF818TKE | ChIP-seq | fold change over control | bigWig |
| *H3K4me1-human* | ENCFF587JLN | ChIP-seq | fold change over control | bigWig |
| *H3K4me1-human* | ENCFF336HIU | ChIP-seq | fold change over control | bigWig |
| *H3K4me1-human* | ENCFF591KWL | ChIP-seq | fold change over control | bigWig |
| *H3K4me1-human* | ENCFF936LEU | ChIP-seq | fold change over control | bigWig |
| *H3K23ac-human* | ENCFF461ICR | ChIP-seq | fold change over control | bigWig |
| *H3K23ac-human* | ENCFF904OAU | ChIP-seq | fold change over control | bigWig |
| *H3K23ac-human* | ENCFF464QEO | ChIP-seq | fold change over control | bigWig |
| *H3K4me3-human* | ENCFF170TVY | ChIP-seq | fold change over control | bigWig |
| *H3K4me3-human* | ENCFF819TZS | ChIP-seq | fold change over control | bigWig |

| | | | | |
|---|---|---|---|---|
| *H3K4me3-human* | ENCFF372YWG | ChIP-seq | fold change over control | bigWig |
| *H4K5ac-human* | ENCFF047DYU | ChIP-seq | fold change over control | bigWig |
| *H4K5ac-human* | ENCFF716DGP | ChIP-seq | fold change over control | bigWig |
| *H4K5ac-human* | ENCFF114DFQ | ChIP-seq | fold change over control | bigWig |
| *H3K9ac-human* | ENCFF978MIP | ChIP-seq | fold change over control | bigWig |
| *H3K9ac-human* | ENCFF238LTR | ChIP-seq | fold change over control | bigWig |
| *H3K9ac-human* | ENCFF504HTE | ChIP-seq | fold change over control | bigWig |
| *EP300-human* | ENCFF020IGM | ChIP-seq | fold change over control | bigWig |
| *EP300-human* | ENCFF211LTW | ChIP-seq | fold change over control | bigWig |
| *EP300-human* | ENCFF491ZOF | ChIP-seq | fold change over control | bigWig |
| *H3K4me3-human* | ENCFF818ROA | ChIP-seq | fold change over control | bigWig |
| *H3K4me3-human* | ENCFF546MMF | ChIP-seq | fold change over control | bigWig |
| *H3K4me3-human* | ENCFF603GRK | ChIP-seq | fold change over control | bigWig |
| *H4K8ac-human* | ENCFF938PJQ | ChIP-seq | fold change over control | bigWig |
| *H4K8ac-human* | ENCFF776JHS | ChIP-seq | fold change over control | bigWig |
| *H4K8ac-human* | ENCFF510WQU | ChIP-seq | fold change over control | bigWig |
| *H3K27me3-human* | ENCFF239OTP | ChIP-seq | fold change over control | bigWig |
| *H3K27me3-human* | ENCFF502GXT | ChIP-seq | fold change over control | bigWig |
| *H3K27me3-human* | ENCFF330TNW | ChIP-seq | fold change over control | bigWig |
| *H4K91ac-human* | ENCFF649HHW | ChIP-seq | fold change over control | bigWig |
| *H4K91ac-human* | ENCFF131DYG | ChIP-seq | fold change over control | bigWig |
| *H4K91ac-human* | ENCFF068LXN | ChIP-seq | fold change over control | bigWig |
| *H2BK15ac-human* | ENCFF152OXY | ChIP-seq | fold change over control | bigWig |
| *H2BK15ac-human* | ENCFF529TKB | ChIP-seq | fold change over control | bigWig |

| | | | | |
|---|---|---|---|---|
| *H2BK15ac-human* | ENCFF236YZE | ChIP-seq | fold change over control | bigWig |
| *H3K79me1-human* | ENCFF944ROK | ChIP-seq | fold change over control | bigWig |
| *H3K79me1-human* | ENCFF465TGW | ChIP-seq | fold change over control | bigWig |
| *H3K79me1-human* | ENCFF349YSW | ChIP-seq | fold change over control | bigWig |
| *H3K18ac-human* | ENCFF649UOB | ChIP-seq | fold change over control | bigWig |
| *H3K18ac-human* | ENCFF413LVW | ChIP-seq | fold change over control | bigWig |
| *H3K18ac-human* | ENCFF592COP | ChIP-seq | fold change over control | bigWig |
